# A time-resolved axillary volatile fingerprint of human fear

**DOI:** 10.64898/2026.05.11.724357

**Authors:** T. Bruderer, A. Greco, M. Ripszam, A.L. Callara, A. Gargano, S. Reale, F.M. Vivaldi, D. Biagini, N. Vanello, M. Perchiazzi, T. Lomonaco, E.P. Scilingo, F. Di Francesco

## Abstract

Human axillary odour can convey information about emotional state, but the molecular composition of fear related volatile emissions remains undefined. Here, we combine low background dual chamber axillary sampling, synchronous real time and offline mass spectrometry, chemical domain group independent component analysis and hierarchical Bayesian modelling to identify a molecular fingerprint of acute fear during immersive virtual reality fear induction. In 37 healthy adults, this framework recovered five axillary volatile components associated with a continuous physiology derived fear index. The fingerprint comprised increased emissions of acetic, butyric, caproic and caprylic acids, octanal and acetone, together with decreased sulcatone, decanal, geranylacetone and citraconic anhydride. This coordinated pattern was reproducible across individuals and emerged from unsupervised chemical decomposition followed by regularised Bayesian selection, without prior biochemical constraints. Posterior predictive checks and leave one participant out refitting supported its robustness within the cohort. Its composition and directionality are consistent with sympathetic autonomic metabolic mobilisation, suggesting contributions from fatty acid handling, acetone related metabolism and axillary gland output. These findings define a candidate molecular cue of acute human fear and establish a replicable methodological template for decoding the chemical language of human emotion.

## Introduction

Chemical communication is the most ancient and phylogenetically widespread channel through which animals signal danger to conspecifics [1–3]. Across taxa as distant as fish, rodents, and ungulates, predator exposure or social stress reliably triggers the release of volatile compounds that elicit defensive behaviours, autonomic activation, and endocrine changes in receivers exposed to the odour alone [1,3–5]. Whether the same form of chemically mediated alarm signalling has been retained in humans, despite our heavy reliance on visual and verbal communication, is a question that has been debated for more than five decades and remains, to date, unresolved [6–8].

Since the first systematic demonstrations that human axillary odour can convey emotional state, behavioural and neuroimaging evidence have provided convergent, albeit indirect, support for the existence of a human fear-related chemosensory cue [9–10]. Sweat collected from donors undergoing acute psychological stress (e.g., parachute jumps, anticipatory speech tasks, threat-related films [10–13]) has been shown to bias receivers towards threat-congruent facial expressions, to enhance startle reflexes, to recruit fear-related neural circuits including the amygdala and the insula, and to modulate risk-taking behaviour [9,11–15]. A meta-analysis confirmed that these effects, though individually modest, are statistically robust across studies and laboratories [9]. Yet despite this consistent receiver-side evidence, the chemical composition of the putative cue has remained elusive, leaving the field unable to progress from demonstration to mechanism [7,16].

Two methodological obstacles help explain this limited progress. The first concerns how samples are typically collected. Most studies still rely on absorbent pads worn under the axilla and later desorbed in the laboratory. This approach can introduce substantial chemical contamination from the pad material itself [17], and it does not provide the temporal resolution needed to follow an evoked emotional response as it unfolds [14,18]. The second obstacle is analytical. Even when high-quality chemical data are available, the methods most commonly used, such as mass-univariate testing of individual features or unsupervised clustering of subjects, are not well matched to the nature of the problem. The relevant signal is likely to be a coordinated multi-compound pattern, shared across individuals but expressed with subject-specific dynamics. At the same time, the predictors are strongly intercorrelated, and the response is a noisy, autocorrelated time series [19–21]. As a result, the literature has produced a fragmented list of candidate compounds which, although individually suggestive and in some cases recurring across laboratories, has not yet converged into a coherent and reproducible molecular fingerprint [7–8,16].

Here we set out to test, with a single integrated framework, the hypothesis that acute human fear is accompanied by a reproducible, biologically interpretable shift in the composition of axillary volatile emissions, i.e., a chemical fingerprint, in the same operational sense in which neuroimaging speaks of functional fingerprints of brain states [22]. Our framework rests on four innovations that together address the bottlenecks above. We developed a 3D-printed, dual-chamber skin sampler manufactured from highly inert polymers that captures volatile organic compounds (VOCs) directly in the gas phase above the axilla, eliminating pad-derived contamination [17]. This enabled, for the same anatomical site and the same emotional episode, both real-time PTR-QTOF monitoring at sub-second temporal resolution and time-integrated offline collection for high-resolution GC×GC-QTOF identification. We elicited acute fear through a previously validated immersive virtual-reality protocol while continuously recording autonomic nervous system correlates [23]. To recover chemical patterns shared across participants, we adapted Group Independent Component Analysis to the chemical domain (CD-ICA), exploiting the formal analogy between voxels in fMRI and mass-spectral channels in PTR-QTOF to extract statistically independent chemical maps with subject-specific time courses [19–20]. Finally, we built a hierarchical Bayesian Elastic-Net regression with first-order autoregressive errors that simultaneously regresses all component time courses against a continuous, physiologically derived fear index, providing principled variable selection and full uncertainty quantification under the dependence structure of the data [21,23–25].

Here, we apply this framework to 37 healthy adults and identify five CD-ICA chemical components associated with a continuous physiological fear index [23]. These components define a reproducible axillary volatile fingerprint of acute fear, dominated by short-and medium-chain fatty acids and acetone, alongside a reduction of selected skin-derived volatiles. The molecular composition and directionality of this fingerprint support an interpretation in which acute fear is accompanied by a coordinated reorganization of axillary emissions, compatible with sympathetic autonomic-metabolic mobilization [26–30]. By linking time-resolved axillary chemistry to a physiological index of fear within a hierarchical Bayesian framework, our study provides a chemically defined candidate cue for human fear chemosignalling and a methodological template for mapping the molecular correlates of other affective and social states [7–9]. The receiver-side signalling, i.e. the possible behavioural and neuroimaging effects of the identified chemical cue, has not been tested.

## Results

### A low-background sampling system for synchronous real-time and cumulative analysis of axillary volatiles

A robust test of the fear-fingerprint hypothesis required resolving the sampling problem which we recently uncovered and that has historically constrained the field [17]. To this aim, a custom 3D-printed dual-chamber skin sampler (Fig. 1a) was designed and manufactured from a polycarbonate blend selected for its very low VOC background, with all internal connections in PTFE and PFA. The two chambers share a single skin-facing aperture but route the collected gas to two parallel analytical pipelines: one chamber delivers the sample, in real time and at sub second-scale temporal resolution, to a high-resolution PTR-QTOF mass spectrometer (Vocus 2R, mass resolving power ≈ 14,000); the other exposes a HiSorb sorbent probe that is subsequently thermally desorbed into a two-dimensional gas chromatograph coupled to a quadrupole time-of-flight mass spectrometer (GC×GC-QTOF). Critically, a pre-mixed gas mixture of 13 VOCs standards (e.g. xylene, acetonitrile, α-pinene, and others) is continuously co-injected through the sampler inlet at a known low gas-phase concentration (i.e., 10 ppb), providing a built-in quality-control trace that allows post-hoc detection of motion artefacts, leaks, and instrumental drift on a per-subject basis.

**Fig. 1.**
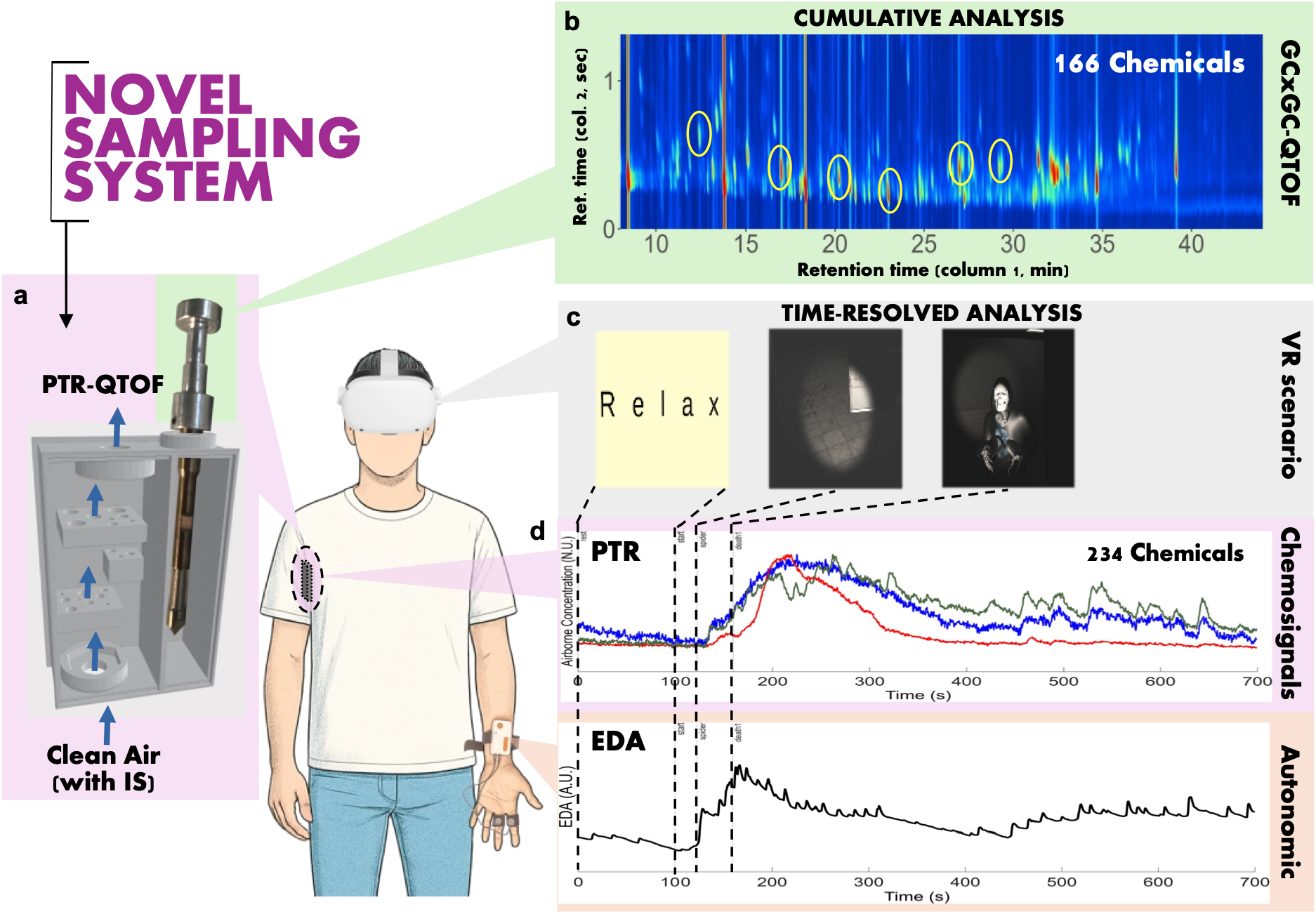
A low-background sampling system for synchronous real-time and cumulative analysis of axillary volatiles. (a) Schematic of the dual-chamber 3D-printed sampler, showing the PTR-QTOF online channel, the continuous internal-standard injection, and the HiSorb offline chamber for subsequent analysis of (b) the cumulative samples with GCxGC-QTOF for compound identification (166 chemicals). (c) Three-phase virtual-reality protocol (resting, neutral living room, and abandoned-hospital fear scenario). (d) Representative single-subject multimodal recording, with concurrent PTR-QTOF chemical traces (234 features), electrodermal activity, and discrete-event timestamps from the VR scenario.

Forty-five healthy young adults (28 female; mean age 23.5 ± 2.4 years; 17 male; mean age 23.7 ± 2.2 years) were recruited under stringent pre-experimental controls (smoking, alcohol, strong-flavoured foods, and physical exercise abstinence in the 24 hours preceding the session, standardised body wash the previous evening, no coffee on the day of measurement). Each participant underwent a previously validated immersive virtual-reality protocol comprising a 5-minute resting baseline, a 5-minute neutral room exploration, and, after a 100-second resting baseline, a 10-minute fear-induction phase set in an abandoned hospital populated with up to 12 fear-eliciting stimuli (jumpscares, predatory animals, supernatural figures), all delivered through an Oculus Rift S head-mounted display while standing [23]. Throughout the session, axillary VOCs, electrodermal activity, and event timestamps were acquired synchronously, generating a multimodal time-resolved dataset for human chemosignal research (Fig. 1b-d). After application of predefined exclusion criteria addressing residual sweat carry-over, leak events confirmed by the internal-standard trace, and an instrumental fault, the final dataset comprised 37 participants (14 males, 23.4 ± 2.3 years; 23 females, 23.9 ± 2.7 years). The virtual reality (VR) fear scenario increased self-reported fear, STAI-Y1 state anxiety, and electrodermal sympathetic activity relative to the neutral scenario. Self-reported fear increased from a median of 1 after the neutral scenario to 5 after fear induction (two-tailed paired Wilcoxon signed-rank test, n = 45, Z = −4.51, p = 6.53 × 10−6, r = 0.67; Hodges–Lehmann paired difference = 2.5, 95% CI [1.5, 3.0]; Supplementary Table 4). STAI-Y1 state anxiety also increased from a median of 37 to 42 (n = 45, Z = −2.16, p = 0.030, r = 0.32; Hodges–Lehmann paired difference = 4.0, 95% CI [0.5, 7.5]; Supplementary Table 4). Consistently, electrodermal sympathetic activity increased during fear relative to neutral stimulation (n = 44, Z = −5.77, p = 8.16 × 10−9, r = 0.87; Hodges–Lehmann paired difference = 5.2, 95% CI [3.6, 6.6]; Supplementary Table 5).

### CD-ICA recovers shared chemical fingerprints with subject-specific dynamics

The PTR-QTOF preprocessing pipeline (background correction, high-resolution mass-axis recalibration, automated peak detection, elemental-composition assignment with isotopic and water-adduct cross-checks, and General Linear Model regression of three robust calibrant traces to remove technical variance) yielded 234 detected m/z features, of which 129 passed the baseline-versus-blank and fear-versus-baseline inclusion filters and were retained for CD-ICA (Supplementary Table 1). To extract chemical patterns expressed consistently across subjects but with individual temporal dynamics, we adapted to the chemical domain the Group Independent Component Analysis framework that has become standard in functional neuroimaging [19]. The resulting Chemical-Domain ICA (CD-ICA) treats mass-spectral channels as the analogue of voxels and feature-time as the analogue of fMRI scan-time and using subject-specific PCA for dimensionality reduction, noise-free ICA, and the GICA3 for back-reconstruction (Fig. 2a). This decomposition rests on two assumptions tailored to chemosignal research: (i) the chemical composition of each independent component—its IC map—is shared across participants, reflecting common underlying physiological processes; and (ii) each subject expresses the components with their own time courses, accommodating the inevitable heterogeneity in how individuals explore and react to the VR environment [23]. Analogous to component-based EEG and fMRI analyses, the extracted ICs likely encompassed both biologically meaningful sources of chemical activity and non-biological contributions, including subject-or sensor-related motion, instrumental noise, and other nonphysiological fluctuations.

**Fig. 2.**
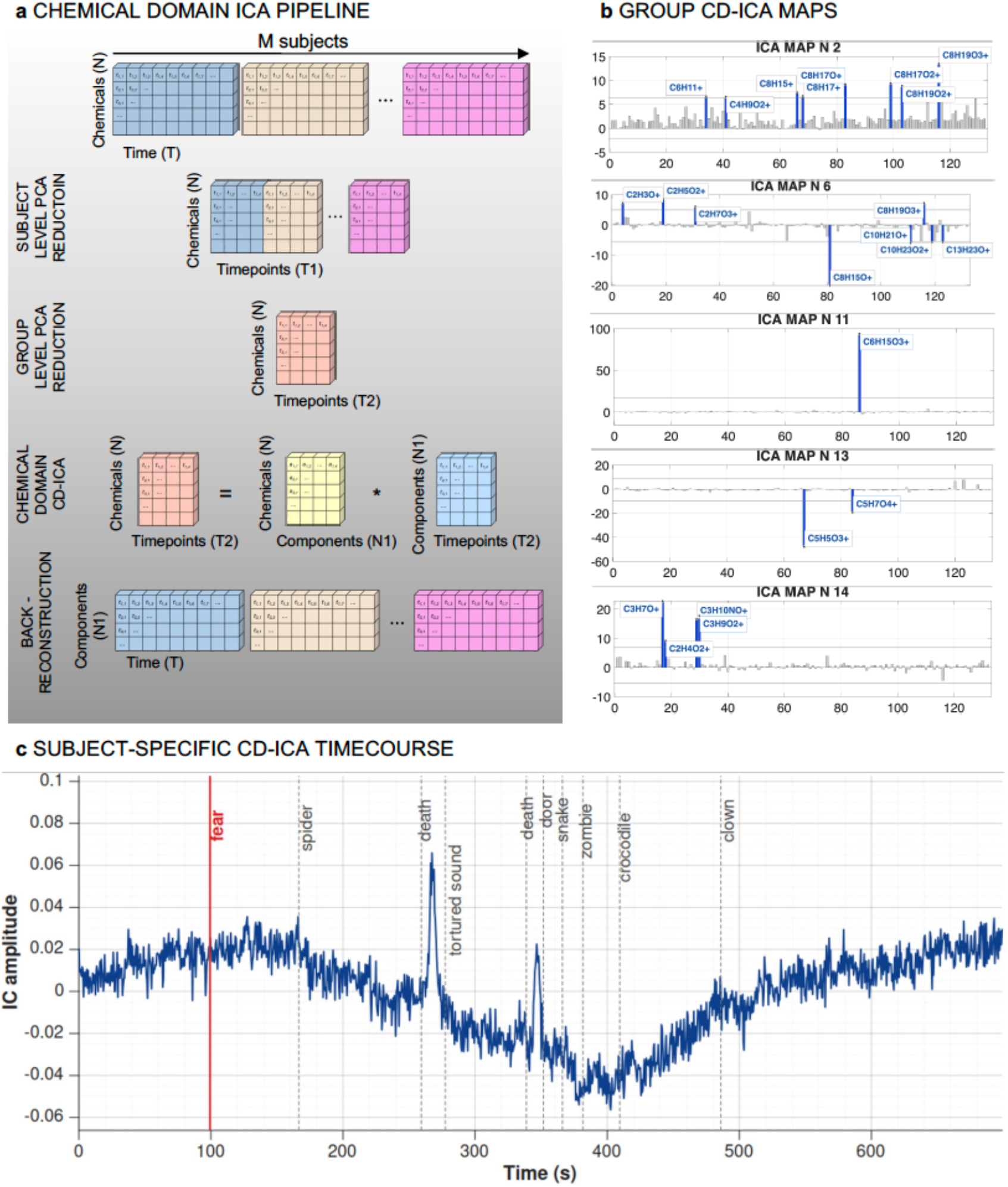
Group Chemical-Domain ICA recovers chemical fingerprints shared across participants. (a) Schematic of the CD-ICA pipeline: subject-specific PCA, temporal concatenation, group-level PCA, noise-free ICA, and GICA3 back-reconstruction yield, for each of 40 components, a shared chemical map, and a set of subject-specific time courses. (b) The five components subsequently identified by the Bayesian model as credibly associated with the fear index (IC2, IC6, IC11, IC13, IC14), shown as chemical loading maps on the m/z axis with sign-corrected loadings. The displayed sign of each peak reflects the product of the IC map weight and the Bayesian regression coefficient, so that positive peaks correspond to compounds emitted more strongly during fear and negative peaks to compounds suppressed during fear. Horizontal dashed lines indicate ±2 standard deviations from the mean loading within each component and are used only to visualise and label the dominant contributing m/z features; statistical selection of fear-associated components was performed by the hierarchical Bayesian model. (c) Representative subject time course for a fear-positive component (i.e., IC2), overlaid with the discrete fear-eliciting events of the VR scenario.

CD-ICA decomposed the group-level dataset into 40 independent components, each defined by a chemical map (the loading of every PTR-QTOF feature on the component) and a set of 37 subject-specific time courses (Fig. 2b,c). The IC maps were chemically interpretable: several components loaded on coherent groups of related ion features, including possible adducts and fragments, consistent with genuine chemical patterns rather than isolated mathematical artefacts [20,31–32]. We note that, because ICA is sign-indeterminate, the biologically meaningful direction of each compound on each component is given by the product of the sign of the loading in the IC map and the sign of the regression coefficient β estimated by the downstream Bayesian model; throughout this paper, we report directionality as that product, so that a compound described as “increasing with fear” is one whose actual axillary emission rises during fear episodes irrespective of the internal sign convention of the decomposition. Crucially, the IC maps were extracted blindly with respect to both the experimental condition and the fear timeline, so that any subsequent association between a component and the fear response constitutes an independent statistical test rather than a circular confirmation.

Before entering the Bayesian regression, the 40 extracted components were subjected to a blind component-quality screening analogous to artefact rejection in EEG and fMRI ICA analyses. Components dominated by non-physiological patterns—isolated noise features, residual washout features, or isolated contaminant features—were excluded from the inferential model. This procedure removed 11 components and retained 29 chemically interpretable CD-ICA components for downstream hierarchical Bayesian modelling.

### A hierarchical Bayesian Elastic-Net model identifies five chemical components credibly associated with the physiological fear index

To identify which of the chemical components were credibly associated with the fear response, we developed a hierarchical Bayesian Elastic-Net regression model with first-order autoregressive errors. The architecture was chosen to address, jointly and within a single principled framework, three statistical features of the data that are typically handled in isolation, if at all, in chemosignal research: the nesting of repeated measurements within subjects (modelled through subject-specific random intercepts drawn from a common population distribution), the serial dependence of consecutive observations along each subject’s time course (modelled as an AR(1) process whose stationary variance correctly initializes the first observation of each subject), and the simultaneous high dimensionality and multicollinearity of the 29 candidate predictors (handled by augmenting the log-posterior with explicit L1 and L2 penalty potentials whose strengths λL1 and λL2 are themselves inferred from the data through Exponential hyperpriors).

Of note, to test the fingerprint hypothesis, the dynamics of the chemical components were regressed against an objective, time-resolved physiological proxy of the participant’s fear physiological state. Accordingly, we derived such a proxy from the electrodermal signal, exploiting the previously validated model that maps the standard deviation of the slow tonic skin-conductance level and the cumulative amplitude of the phasic sudomotor-nerve burst activity onto a self-reported subjective fear scale [23]. Applying this model to non-overlapping 5-second windows along the fear-induction phase yielded, for each participant, a continuous fear index sampled at the same temporal resolution as the chemical components (Fig. 3a).

**Fig. 3.**
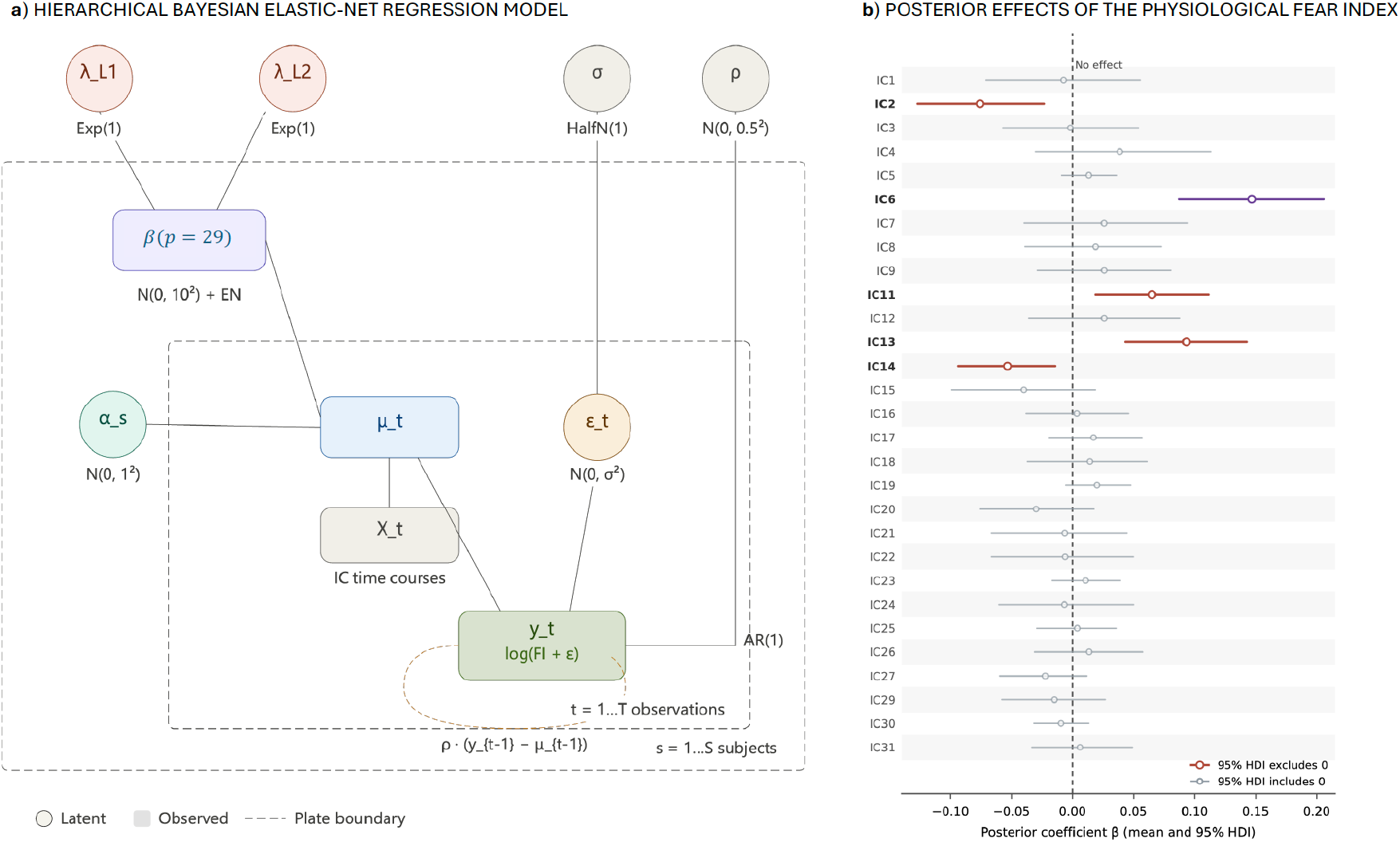
A hierarchical Bayesian Elastic-Net model with AR(1) errors links chemical fingerprints to the physiological fear index. (a) Plate diagram of the generative model, showing subject-specific intercepts, the AR(1) error process, the Elastic-Net regularised coefficient vector β, and the data-driven Exponential hyperpriors on the L1 and L2 penalty strengths. (b) Forest plot of the posterior distributions of all β coefficients (mean and 95% HDI), with the five components whose 95% HDI excludes zero highlighted (IC2, IC6, IC11, IC13, IC14). The dashed vertical line marks the no-effect value (β = 0).

Because the fear index is strictly positive and right-skewed, we modelled its log-transform under a Gaussian likelihood, ensuring that posterior predictive samples respect the support of the response. Posterior inference was performed using NUTS (4 chains, 4,000 iterations each, 2,000 discarded as warm-up; R̂ = 1.00 for all parameters; effective sample size > 6,000 for every parameter of interest, see Table 1).

**Table 1.** Posterior summaries for the five chemical components credibly associated with the fear index. Posterior mean and standard deviation, 95% Highest Density Interval (HDI), bulk effective sample size (ESS), and Gelman–Rubin diagnostic (R̂) [25] for the regression coefficient β of each credibly-associated component. The final column reports the biologically meaningful direction of the dominant compounds in each component (i.e., the product of the sign of β and the sign of the loading in the IC map; see text). All five components achieve R̂ = 1.00 and ESS > 6,000, indicating excellent MCMC convergence and posterior sampling efficiency [24–25].

| Component | Posterior mean ($\beta$) | Posterior SD | 95% HDI | ESS bulk | $\hat{R}$ | Direction vs. fear |
| --- | --- | --- | --- | --- | --- | --- |
| IC2 | -0.076 | 0.027 | [-0.127, -0.023] | 12,264 | 1.00 | positive (↑) |
| IC6 | +0.147 | 0.030 | [+0.087, +0.205] | 12,502 | 1.00 | mixed |
| IC11 | +0.065 | 0.024 | [+0.019, +0.111] | 10,275 | 1.00 | positive (↑) |
| IC13 | +0.093 | 0.025 | [+0.043, +0.142] | 14,134 | 1.00 | negative (↓) |
| IC14 | -0.053 | 0.020 | [-0.094, -0.014] | 12,816 | 1.00 | positive (↑) |

A component was deemed credibly associated with fear if the 95% Highest Density Interval (HDI) of the posterior distribution of its regression coefficient β excluded zero. Five components met this criterion (Fig. 3b; Table 1): IC2 (β = −0.076; 95% HDI [−0.127, −0.023]), IC6 (β = +0.147; [+0.087, +0.205]), IC11 (β = +0.065; [+0.019, +0.111]), IC13 (β = +0.093; [+0.043, +0.142]) and IC14 (β = −0.053; [−0.094, −0.014]). The remaining 24 retained components had β posteriors centered on, or close to, zero, suggesting that the Elastic-Net penalty regularised weakly informative chemical patterns towards zero and that the surviving components reflect fear-related structure rather than overfitting to noise. The strongest single association was carried by IC6, whose posterior mean coefficient was approximately twice the magnitude of the next-largest credible component, identifying it as the dominant statistical anchor of the fingerprint. Once the loading-by-coefficient sign products are taken into account, four of the five components describe compounds whose axillary emission rises with fear, and one component (IC13) describes a compound whose emission falls with fear; IC6 carries both directions on different feature subsets, in a manner that turns out to be informative about the underlying physiology (see next section).

### The fear fingerprint converges on sympathetic metabolic reorganisation

We next identified the molecular constituents of the five fear-associated components by combining the high-resolution elemental-formula assignment provided by PTR-QTOF with the isomer-resolving and NIST library-matching capacity of the offline GC×GC-QTOF analysis of HiSorb-collected sweat. The latter, performed on the same anatomical site and during the same experimental session on pooled fear and blank samples, retained 166 compounds out of 296 detected chromatographic features after FDR-corrected blank subtraction (fold-change > 3, adjusted p < 0.01), and a subsequent volcano-plot comparison between fear and neutral conditions (fold-change > 1.5, adjusted p < 0.05, FDR-corrected). The composition and directionality of each of the five fear-associated CD-ICA components, derived from the sign-corrected loadings on the IC maps, are summarised in Table 2. All fear-modulated chemical components could either be identified by direct match with the 166 identified compounds or by specific mass differences related to water adducts, water losses, or ammonium adducts and their high correlation across the time-domain with one of the identified compounds. Identifications should be regarded as high-confidence putative assignments (Supplementary Tables 2-3).

**Table 2.**
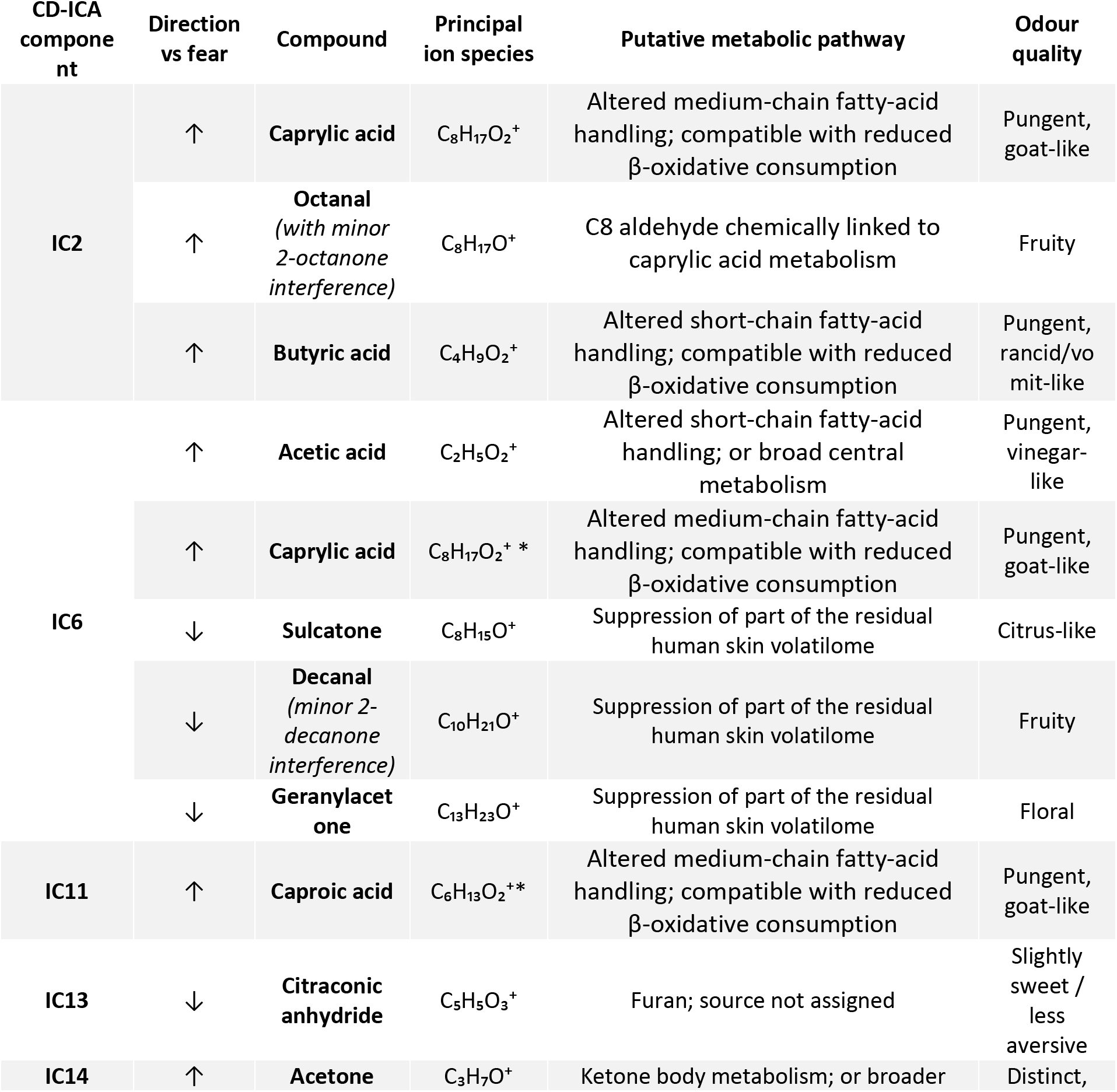
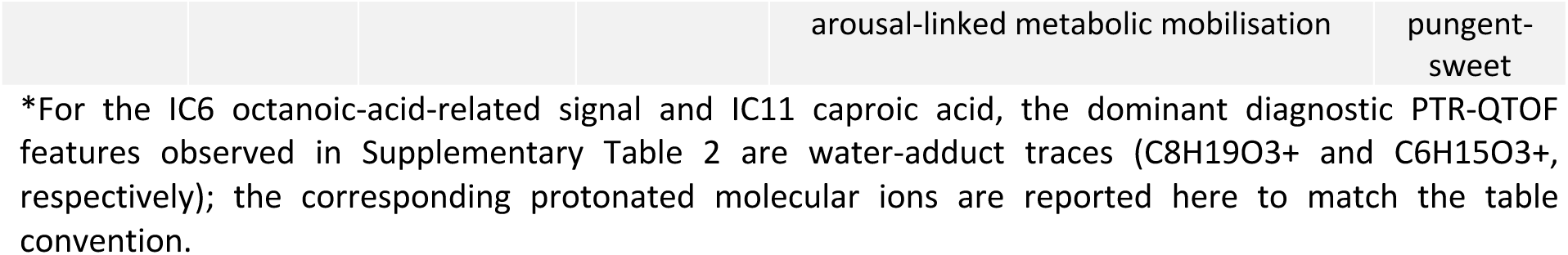
The human fear chemical fingerprint. . Molecular composition of the five CD-ICA components credibly associated with the physiological fear index (95% HDI of β excluding zero), with sign-corrected directionality (↑ increase, ↓ decrease with fear), putative metabolic pathway, and odour quality. Ion species refers to the principal protonated molecular ion; full ion characterisation, including water adducts, water-loss fragments, and ammonia adducts, is provided in Supplementary Table 2. Identification was achieved by combined real-time PTR-QTOF elemental-formula assignment and offline GC×GC-QTOF library matching.

A coherent and biologically constrained molecular picture emerges. IC2 carries a short-and medium-chain fatty-acid block comprising butyric acid, caprylic acid, the C8 aldehyde octanal, all increasing with fear. Caprylic acid was identified by its protonated ion C8H17O2+ with GC×GC-QTOF library support which was highly correlated to its water adduct C8H19O3+, Octanal, a C8 aldehyde chemically linked to caprylic acid metabolism [33–34], co-loaded on the same component through its protonated ion C8H17O+ and was highly correlated with its water adduct C8H19O2+, while GC×GC-QTOF indicated a minor interference of the isomer 2-octanone which cannot be distinguished with PTR-QTOF alone. Butyric acid was identified within the same component through its protonated ion C4H9O2+ and confirmed by GC×GC-QTOF library matching [35–36]. IC6, the strongest single component of the fingerprint, contains within a single chemical pattern two physiologically coherent but oppositely directed contributions: acetic acid increases with fear, identified by its protonated ion C2H5O2+ alongside its highly correlated water-loss fragment C2H3O+ and water adduct C2H7O3+, together with the water adduct of caprylic acid C8H19O3+ that is highly correlated with the corresponding protonated caprylic acid ion. By contrast, three skin-resident volatiles decrease with fear in the same component: sulcatone (6-methyl-5-hepten-2-one, ion C8H15O+), decanal (ion C10H21O+, with a minor interference of the 2-decanone water adduct ion C10H23O2+), and geranylacetone (6,10-dimethyl-5,9-undecadien-2-one, identified through C13H23O+) [37–39]. IC11 is dominated by caproic acid, identified through its water adduct C6H15O3+ and assigned to the corresponding protonated ion of caproic acid; this compound increases with fear. IC13 is dominated by the furan citraconic anhydride, respectively 3-methyl-2,5-furandione (ion C5H5O3+, with water adduct C5H7O4+ providing cross-confirmation); this compound decreases with fear. Finally, IC14 is dominated by acetone, identified by a four-ion pattern—its protonated molecular ion C3H7O+, and three highly correlated ions, its water adduct C3H9O2+, its ammonium adduct C3H10NO+, and a likely instrument artefact peak C2H4O2+ related to the high acetone levels close to saturation levels —all converging on the same molecule, which increases with fear [37,40].

Considered together, the constituents of the fear fingerprint converge on a biologically coherent reading. The synchronous rise of four short-and medium-chain volatile fatty acids—acetic acid in IC6, butyric and caprylic acids in IC2, and caproic acid in IC11—is consistent with a fear-associated shift in short-and medium-chain fatty-acid handling, potentially reflecting reduced β-oxidative consumption under acute sympathetic arousal [27–28,40]. This interpretation is compatible with the well-documented metabolic reorientation from mitochondrial fatty-acid β-oxidation towards faster carbohydrate-based energy mobilisation during acute stress [27–29]. Together with the parallel increase in acetone, a volatile end-product associated with an alternative energy supply through ketone-body-related metabolism, this pattern points to a coordinated fear-associated metabolic mobilisation, in which altered fatty-acid handling and ketone-body-related chemistry become detectable in the axillary headspace [37,40]. The resulting volatile profile is therefore best interpreted as a peripheral chemical readout compatible with the sympathetic energy demands of an acute defensive response [28–29]. Octanal in IC2, the corresponding C8 aldehyde chemically linked to caprylic acid metabolism, fits the same metabolic block [33–34]. The decrease of the three skin-derived volatiles (sulcatone, decanal with a minor 2-decanone contribution, and geranylacetone in IC6) and of the furan citraconic anhydride in IC13 is most parsimoniously interpreted not as a metabolic effect but as a consequence of the abrupt change in axillary sweat-gland output that accompanies sympathetic activation, transiently diluting or displacing the resident skin volatilome [37] [38–39].

The structure of IC6 is particularly informative in this respect. That a single statistically independent component should carry, simultaneously, a positive loading on acetic acid and negative loadings on three skin-resident volatile signals is consistent with the physiological model we propose: a single sympathetic arousal state may produce both the metabolic increase in volatile fatty acids and the suppression of resident skin metabolites, and these two effects are therefore expected to vary across time and across subjects. The recovery of this anti-correlated pattern as a single independent component, rather than as two separate components, is therefore not a quirk of the decomposition but a fingerprint of the unified physiological process that generates both effects. Crucially, this metabolic reading was not built into the model: the IC maps were extracted blindly by an unsupervised decomposition, and the regression coefficients were selected by a regularised Bayesian model that had no access to the underlying biochemistry. The convergence of an unsupervised chemical decomposition, a blind statistical selection, and a coherent biochemical interpretation thus constitute mutually independent lines of evidence that the fingerprint we describe is compatible with an acute, sympathetically linked metabolic reorganisation [40] [27–29].

It is also notable that the affective valence of the fingerprint’s odour profile is internally consistent with an alarm-signalling function [1,9]. The compounds that increase with fear are dominated by pungent, aversive odours—butyric acid, the prototypical reference odour of vomit; caproic and caprylic acid, both pungent and goat-like; acetic acid, the sharp odour of vinegar; and acetone, with its distinctive and pungent quality [37,41–44]. The single fruity exception in the rising group, octanal, is chemically linked to caprylic acid metabolism and is unlikely to dominate the perceptual mixture [33–34,43]. The compounds that decrease with fear, conversely, carry pleasant qualities (citrus and floral for sulcatone and geranylacetone, fruity for decanal, slightly sweet for citraconic anhydride) [43]. The net olfactory consequence of the fear-evoked volatile reorganisation is therefore a shift of the axillary headspace simultaneously towards an aversive odour profile and away from a pleasant one—a property compatible with a chemosignalling function that we did not optimise for in our analysis but that emerges naturally from the molecular composition and directionality of the fingerprint [7–9,11–15]. This interpretation is consistent with a recent cross-cultural body-odour lexicon in which stress body odours were frequently described as sour, unpleasant, strong, and pungent [45].

### Robustness of the fear fingerprint

Two additional analyses supported the robustness of the fear fingerprint to model misspecification and single-participant influence. First, posterior predictive checks showed that datasets simulated from the fitted hierarchical Bayesian Elastic-Net model reproduced the main distributional and temporal properties of the observed response. The observed mean of the log-fear-index (−1.25) fell within the 95% posterior predictive interval [−1.29, −1.18] and was close to the posterior predictive mean (−1.23; two-sided PPC p = 0.536). The same was true for the variance (observed = 1.18; posterior predictive mean = 1.17; 95% interval [1.09, 1.27]; p = 0.924) and the lag-1 autocorrelation (observed = 0.488; posterior predictive mean = 0.498; 95% interval [0.465, 0.531]; p = 0.540), indicating that the model captured both the scale and temporal dependence of the physiological response. Second, we re-fitted the model in a leave-one-subject-out scheme across the 37 participants with valid EDA-derived fear-index time series. Four of the five fear-associated components remained credible in all 37 folds (IC6, IC11, IC13 and IC14), while IC2 remained credible in 36 of 37 folds. Posterior means were stable across folds (IC2, β = −0.0748 ± 0.0101; IC6, β = 0.1477 ± 0.0071; IC11, β = 0.0645 ± 0.0054; IC13, β = 0.0926 ± 0.0074; IC14, β = −0.0534 ± 0.0049), with no systematic shrinkage towards zero. These analyses indicate that the fingerprint is not driven by a single influential participant and that the Bayesian AR(1) model adequately captures the distributional and temporal structure of the fear-index data.

## Discussion

For more than half a century, the question of whether human emotions can be transmitted between individuals through volatile chemical cues has occupied an uncomfortable middle ground between behavioural plausibility and molecular obscurity [6–8]. Receivers exposed to fear sweat respond with measurable shifts in perception, neural activation and behaviour [9,11–15], and yet the chemical content of that sweat—the very thing that one would have to identify, characterise and ultimately synthesise to close the explanatory loop—has remained, until now, a fragmented set of partially overlapping candidates whose organisation into a reproducible molecular fingerprint has remained unresolved [7,16]. The work we present here changes the terms of this question. By coupling a low-background, time-resolved sampling system to an analytical framework explicitly designed to recover patterns shared across individuals while quantifying their relationship to a continuous physiological readout of emotion, we identify a chemical fingerprint of acute human fear that is reproducible, biologically interpretable, and statistically credible under stringent and multiply independent criteria. The fingerprint is not an arbitrary list of compounds: its molecular composition—four short-and medium-chain volatile fatty acids, the C8 aldehyde octanal, and acetone, all rising synchronously with fear, paired with the suppression of three skin-derived ketones and a furan—maps onto a well-characterised pair of metabolic pathways that the sympathetic nervous system is known to engage during acute stress [26–30,37,40]. We therefore advance a mechanistic hypothesis: the fear-associated fingerprint may reflect the volatile readout of an integrated defensive state emerging from the coupling between central threat processing, interoceptive feedback and peripheral autonomic-metabolic mobilisation [7–9,26–30,38–39,46].

The central inference enabled by our results is the articulation of a mechanistic link that, in human chemosignal research, has not previously been specified end-to-end [7–8]. An external threat—here delivered through a controlled virtual-reality scenario—is expected to engage central threat-processing circuits together with peripheral autonomic pathways, including the sympathetic branch of the autonomic nervous system [23,28–29,46]. Sympathetic activation is tightly coupled to systemic metabolic mobilisation, including coordinated regulation of glucose and lipid metabolism and engagement of ketone body metabolism pathways under energetic challenge [26–30,40]. Both pathways generate volatile end-products that are sufficiently small, lipophilic and water-soluble to partition into eccrine and apocrine sweat and to evaporate from the skin surface within the timescale of an acute affective episode [18,37–39,47]. The compounds we identify—four short-and medium-chain fatty acids and acetone—are therefore consistent with volatile end-products expected from altered fatty-acid handling and ketone-body-related metabolism under acute sympathetic arousal [26–30,37–40]. The structure of the fingerprint refines this prediction in a way that we did not anticipate but that the data demand: rather than appearing as a uniform increase in fatty acids and acetone superimposed on an otherwise unchanged background volatilome, the fingerprint takes the form of a coupled rise of metabolic end-products and a coupled fall of three resident skin volatiles, with one of the components (IC6) carrying both directions of effect within a single statistically independent pattern. This pattern is consistent with a coordinated physiological reorganisation rather than a passive accumulation of metabolites: the same sympathetic arousal state that mobilises energy metabolism can also alter eccrine and apocrine gland output, making the two effects detectable in the axillary headspace as one coherent event [28–29,47–49]. This convergence across unsupervised decomposition, Bayesian selection, and chemical annotation argues against a purely artefactual origin of the fingerprint. We therefore propose that the human fear fingerprint should be understood, mechanistically, as a peripheral chemical correlate of an integrated defensive state, in which central appraisal, interoceptive feedback, and sympathetically mediated metabolic mobilisation are dynamically coupled. This formulation reframes the problem of human fear chemosignalling as a quantifiable problem in metabolic physiology rather than a contested phenomenon in pheromone biology, and it predicts that the fingerprint will be modulated, in lawful and testable ways, by any intervention that alters sympathetic tone, fatty-acid availability, or eccrine-gland activity [7,16,27–30,38–39,47].

At the same time, this interpretation should be read as a mechanistic hypothesis rather than direct proof of pathway switching. The volatile products we measure show that the relevant carbon-chain classes are detectable in axillary headspace, and clinical and experimental assays in skin-derived cells demonstrate that fatty-acid oxidation and pyruvate/glucose-oxidation pathways can be quantified in human fibroblasts [50–52]. However, our experiment did not measure intracellular fluxes, β-hydroxybutyrate, acetoacetate, or glucose/TCA intermediates, and it therefore cannot determine whether fear alters β-oxidation or ketone-body-related pathways within seconds, minutes, or through slower local or systemic processes. Resolving this question will require targeted multi-matrix measurements, ideally combining axillary headspace with blood and skin-surface sweat metabolomics with methods capable of detecting more polar metabolites such as β-hydroxybutyrate, acetoacetate, and glycolytic/TCA intermediates [53–55].

Our results also invite a deliberate clarification of the longstanding debate over whether humans possess pheromones in the strict sense of the term [7,16]. A pheromone, as historically defined in chemical ecology, is an evolved chemical signal selected for its capacity to convey information between conspecifics; a cue, by contrast, is a chemical correlate of an internal state that may or may not have been selected for communicative function [7,16]. The fingerprint we describe is, on the present evidence, a cue: a reliable, mechanistically grounded chemical correlate of an acute emotional state. Whether it has additionally been shaped by selection into a signal—whether, in other words, conspecifics have evolved sensory and neural machinery tuned to its detection and decoding—is a separate empirical question that our study does not attempt to settle [7,16]. By providing a robust molecular description of the cue, our work allows the signal question to be posed in its proper, falsifiable form: do receivers respond, behaviourally and neurally, to the specific molecular composition we identify, in the proportions and at the concentrations at which it is naturally emitted [9]? The closure of this question requires the chemical synthesis of the fingerprint and its delivery to naive receivers under controlled olfactometric conditions, i.e., an experiment that our results render formally specifiable [9,42,56–57]. It is worth noting in this respect that the affective valence of the fingerprint is, on its face, congruent with an alarm-signalling function: the compounds that rise with fear are dominated by pungent, aversive odours classically associated with avoidance behaviour, and the compounds that fall are dominated by pleasant, attractive odours [1,3,9,41–43]. The directionality of the fingerprint thus shifts the perceived odour of the donor towards an aversive profile precisely when the donor is in an aversive emotional state. Whether this olfactory congruence reflects a coincidence of metabolism or a target of natural selection is, again, an empirical question that the synthesis-and-delivery experiment is now positioned to answer [7,16]. Until that experiment is performed, we recommend that the field describe these compounds, and any analogous fingerprints recovered for other emotions, with the chemically and evolutionarily neutral term cue, reserving signal for compounds whose receiver function has been independently established [7,16].

Beyond this specific fingerprint, our framework provides a methodological template for studying human chemical communication at the molecular level, adaptable to other affective states, social contexts, and clinical conditions [7–8]. Each of its three components addresses a problem that has, until now, constrained the field independently [7–8,17]. Low-background dual-chamber sampling resolves the long-standing tension between the analytical advantages of offline mass spectrometry and the temporal resolution required to track an evoked response by performing both simultaneously on the same anatomical site and the same emotional episode [17]. CD-ICA generalises to chemical data the group-decomposition logic that transformed functional neuroimaging from a single-subject discipline into a population-level enterprise and provides a principled solution to the fundamental tension between the assumption that emotions produce conserved chemical patterns and the observation that individuals express those patterns with idiosyncratic dynamics. Importantly, the structure recovered in IC6—a single component carrying coupled, oppositely-directed contributions from metabolically related compounds—demonstrates that the method recovers physiologically meaningful processes rather than merely chemically clustered features, a capacity that mass-univariate or correlation-based approaches structurally lack [19–22]. The hierarchical Bayesian Elastic-Net with AR(1) errors offers, finally, a single coherent statistical engine that absorbs the nesting of repeated measurements, the autocorrelation of physiological time series, and the multicollinearity of high-dimensional volatilome data, providing full uncertainty quantification rather than the binary significance tests that have dominated the literature. Each of these innovations is, in principle, modular: alternative emotion-induction protocols can replace the virtual-reality scenario, alternative physiological readouts (heart rate variability, pupillometry, EEG correlates of arousal, respiratory dynamics) can replace the electrodermal fear index [23,48–49,58–59], and alternative analytical platforms (wearable electronic sensors for chemical or chemical class specific skin volatiles, secondary electrospray ionisation, selected-ion flow-tube mass spectrometry) can replace the PTR-QTOF front end [53–54]. We therefore expect this framework to enable a systematic chemical mapping of the human emotional volatilome (disgust, sexual arousal, contentment, social anxiety, parental bonding, grief) that has, to our knowledge, never been attempted under a unified methodology [8,37].

The implications of these findings extend beyond basic chemical ecology into translational and clinical territory. Three lines of inquiry seem to us particularly natural. First, the fact that an acute fear state produces a quantifiable, time-resolved chemical fingerprint opens the possibility of objective, contact-free monitoring of affective state in clinical populations whose self-report is impaired or unreliable—children with autism spectrum conditions, adults with severe anxiety or post-traumatic stress disorder, individuals in acute psychiatric care, patients in disorders of consciousness—complementing, rather than replacing, the established physiological and behavioural readouts on which these populations are currently assessed [58–59]. Second, the metabolic interpretation of the fingerprint generates falsifiable predictions for clinical conditions characterised by chronic dysregulation of sympathetic tone or of fatty-acid metabolism: panic disorder, generalised anxiety disorder, chronic stress states and burnout should, on our framework, be associated with measurable shifts in the resting axillary volatilome that recapitulate, in attenuated form, the acute fingerprint we describe; conversely, conditions associated with sympathetic blunting (such as certain forms of depression with psychomotor retardation) should produce inverse shifts. Third, and more speculatively, the convergence of the fingerprint on the volatile end-products of β-oxidation and ketogenesis suggests that the same compound classes monitored here in axillary headspace should also be detectable in blood, urine, saliva, and breath, opening the possibility of multi-matrix biomarker panels for affective and stress-related disorders that exploit the redundancy across compartments to improve specificity [37,53–55]. The translation of any of these possibilities into clinical practice will require longitudinal studies in the relevant patient populations, which the modular structure of our framework places within technical reach.

Several limitations should be considered, each of which defines a clear next step for this line of work. The fear state was elicited by a virtual-reality protocol previously validated against subjective and physiological criteria [23]; whether the same fingerprint is recovered in response to ecologically more naturalistic threats (e.g., public speaking, parachute jumps, exposure-therapy protocols, real-world traumatic events) remains to be tested, and the answer to that question will determine whether the fingerprint is specific to the affective state or to its mode of induction [9]. Our cohort, although well-powered for an intensive multi-modal protocol of this kind, is demographically homogeneous (young, healthy, predominantly European); whether the fingerprint generalises across age, sex, ancestry, dietary habits and metabolic phenotypes is an empirical question that larger and more diverse cohorts are now needed to settle, particularly in light of the metabolic interpretation we propose, which predicts modulation by any factor that alters basal fatty-acid availability. The fear index against which we regressed the chemical components, although derived from a previously validated electrodermal model [23], captures one autonomic axis of the fear response; jointly modelling other autonomic and central nervous system correlates would be expected to refine the temporal ground-truth precision with which the chemical fingerprint can be tracked, and to disentangle the contributions of metabolic reorganisation, gland activation and respiratory dynamics that the present design conflates. Finally, our work establishes the donor-side chemical characterisation of the candidate cue but does not test receiver-side responses: i.e., whether naive individuals, exposed to the synthetic fingerprint at physiologically relevant concentrations and proportions, exhibit the perceptual, neural, and behavioural responses that the existing meta-analytic literature would predict [9]. Each of these limitations corresponds to an experiment that our framework now makes formally specifiable, and we intend the present study to be read as a starting point for a programme of receiver-side and mechanistic experiments.

The question of whether humans communicate emotions chemically has, since its formulation, occupied a peculiar epistemic position: behaviourally indicated, neurally plausible, evolutionarily expected, and yet molecularly unspecified [6–9]. By identifying a reproducible chemical fingerprint of acute fear and embedding it within a mechanistic interpretation centred on the coupling between central threat processing, interoceptive feedback, and peripheral autonomic-metabolic mobilisation, we move this question from the domain of contested phenomenology to that of quantitative biology [7–8]. Our study provides a molecular starting point for testing whether, and how, the human nervous system responds to fear-associated volatile cues.

## Methods

### Participants

A total of 45 healthy subjects were recruited (28 female, mean age 23.5 ± 2.4 years; 17 male, mean age 23.7 ± 2.2 years). Participants were pre-screened with an online questionnaire to ensure they were in good health and were nonsmokers, and to exclude those with cardiovascular conditions, physical dysfunction, or mental illness. Starting the day before the measurement, participants were required to abstain from strong-flavoured foods (including strong spices and garlic), alcohol, smoking, recreational drugs, and excessive physical exercise, as well as sexual activity on the morning of the experiment. Participants were instructed to shave the axillary region and shower using a shampoo and body wash provided by the experimenters (Sensinol, Pierre Fabre, Boulogne, France) the evening before. On the measurement day, they were asked to consume no coffee, to wear a tank top or bra allowing easy access to the axilla for sweat sampling, and to refrain from eating two hours beforehand, consuming only water.

The study was approved by the local ethics committee (n. 14/2019). All participants provided written informed consent in accordance with the Declaration of Helsinki. All experiments were performed in research laboratory 116 of the Department of Chemistry and Industrial Chemistry (DCCI) of the University of Pisa, a facility equipped for VOC analysis and comprising a dedicated area for experiments with human participants.

### Experimental design and protocol

#### General procedure

Upon arrival at the laboratory, participants were familiarised with the experimental environment and briefed on the key stages of the protocol. They were informed that axillary monitoring would be conducted throughout the experiment using a sampling system for real-time monitoring of axillary sweat volatiles (see below), that physiological signals would be monitored via wearable sensors, and that emotional stimulation would be delivered through a VR headset. Participants were explicitly notified of their right to withdraw at any time. Following the provision of information regarding data processing and privacy, baseline psychometric questionnaires were administered. The experimental protocol comprised four sequential phases, throughout which the VR headset was worn: a 5-minute resting baseline, a 5-minute neutral stimulation phase, a 100-second resting baseline, and a 10-minute fear-induction phase. Participants completed assessments of their emotional state and perceived anxiety immediately after each phase (see Subjective assessments below). Upon conclusion, a comprehensive debriefing was conducted to disclose the full scope and objectives of the research.

#### Emotional stimulation

The primary goal of the virtual-reality environments was to elicit authentic psychological states of fear, mirroring those that participants might experience in real-world situations. The design philosophy, detailed in our previous work [23], prioritised creating an immersive setup that could effectively induce targeted emotional responses. All scenarios were designed from a first-person perspective to maximise immersion and ecological validity. Movement speed was limited to mitigate cybersickness [60], and conditions of active engagement through animated virtual contents allowed direct interaction between the user and the environment [23].

The fearful scenario was crafted based on established principles from horror media, designed to trigger two primary fear mechanisms [23,61]: fear of the unknown and jumpscares. The setting, an abandoned hospital, engaged both visual and auditory sensory channels [62] through flickering low-intensity lighting and suspenseful background audio. Timed high-volume sound cues were designed to maximise their startling effect. Participants navigated the dark space using a virtual flashlight, equipped with a joystick in their dominant hand, enabling both movement and flashlight direction. The environment was populated with up to 12 discrete fear-eliciting stimuli known for their high fear-elicitation capabilities [23]: flying bats, a crocodile, snakes, spiders, a loud clown picture, a crackling door sound, metal chains rattling, two zombies, a scary nurse, and two Grim Reaper figures.

The relaxation and neutral scenarios were designed to minimise emotional arousal. The relaxation environment consisted of a minimalist blank scene with warm colours and a simple relaxation instruction. The neutral scenario was a virtual living room in which participants could navigate freely; lighting and object colours were carefully controlled to avoid stimulating an emotional response [23].

Each participant was equipped with an Oculus Rift S head-mounted display (Lenovo Technologies and Facebook Technologies, USA; 2560×1440 LCD, 80 Hz refresh rate, 6 degrees-of-freedom tracking, integrated headphones). The HMD was wired to a PC for rendering. Before the experimental session, participants completed a brief introductory scenario in which they followed a predefined path, allowing them to familiarize themselves with navigation and movement within the virtual environment. Participants stood for the entire duration to reduce motion-sickness risks [60]. External stimuli were minimised to intensify presence [63]. The three environments were developed in Unity3D (2019.4.8f1, Unity Technologies, USA), optimised for 78–80 Hz frame rate and 3296×1776 rendering resolution. Audio was delivered in stereo. The application managed the complete experimental protocol, including event timestamps for subsequent data analysis [23].

#### Subjective assessments

Subjective psychological states were quantified through standardised questionnaires administered via a computer-based platform. Prior to the experimental induction, participants completed the Italian version of the State-Trait Anxiety Inventory (STAI-Y2) [64] to establish a baseline for trait anxiety. After each experimental condition (resting baseline, neutral, and fear induction), participants completed the STAI-Y1 [64] to assess state anxiety, alongside the assessment of fear using a 9-point Likert scale (1: “not at all”; 9: “extremely”). Following the final VR exposure, the Cybersickness in Virtual Reality Questionnaire [65] was administered to monitor potential adverse effects. No relevant effects have been reported.

A critical component of the subjective assessment involved self-assessed subjective fear (SASF) levels. Beyond evaluating emotional states perceived throughout the overall scenario, participants provided intensity ratings for each specific fear-eliciting stimulus encountered within the VR environment, using a 7-point Likert scale (1: “not at all scared”; 7: “extremely scared”; 0: absence of perception of the stimulus). These event-specific data points served as the subjective ground truth for the fear index model (see Fear Index Estimation below), enabling a high-resolution mapping of physiological responses to discrete fear-induced events.

### Data acquisition

#### Sampler Description

We developed a novel sampling system for real-time monitoring of sweat volatiles with simultaneous sweat collection for compound identification. We used highly inert materials for the sampler, which included a polycarbonate blend (Prusa Polymers a.s., Czech Republic) for 3D-printing of the sampler, Teflon connectors, and perfluoroalkoxy (PFA) tubing. The 3D-printed sampler contained two sampling chambers. The first chamber was connected to the PTR-QTOF for real-time monitoring of chemical concentrations. The second chamber was equipped with HighSorb sorbent sticks and subsequently analysed by GC×GC-QTOF for compound identification. The custom sampler was optimized with regard to sampler materials, dimensions, and geometry, skin-sampler interface, sampler tubing and connections, gas supply and flow control, turbulent flow mixing elements, controlled temperatures, cleaning of the sampler, and continuous addition of internal standards across the sampling runs.

### Sampler dimensions and geometry

The aim was to maximise the skin sampling area while keeping the sampler volume minimal. The sampler size was defined to account for the smaller armpits of the female participants while also being suitable for the male participants (based on a pre-pilot study with 23 participants, including 6 males, and 17 females). This resulted in the final sampler dimensions of 4.5 cm length, 3 cm width and 1.5 cm height (with 2 cm width for the first chamber and 1 cm width for the second chamber). The sampler was developed for armpit sampling (with highest abundance of apocrine glands) but can be used to sample most skin surfaces on the human body (including the palms with mainly eccrine glands an the upper back (with mainly sebaceous glands). The sampler geometry is shown in Figure 1 which includes the first sampling chamber with turbulent flow mixing elements and Teflon connectors for real-time monitoring, and the second chamber for integrative sample collection with an insert for the HighSorb samplers.

### Skin-sampler interface

The sampler had to be leak tight which was finally achieved by using velcro straps with a suitable connector and guide as part of the 3D-printed sampler structure. This solution also guaranteed sufficient comfort on-body sampler wearability for sampling periods up to 60 minutes.

### Cleaning of the sampler tubing and connections

We used highly inert perfluoro alkoxy alkane (PFA) tubing (0.8 mm in diameter, 1.5 m in length). The sampler connections had to be from materials that do not result in cold spots (i.e., avoid stainless steel) and are readily cleanable. We used custom-made Teflon connectors for this purpose.

### Gas supply and flow control

We connected the clean air supply from the PTR instrument to the sampler. The clean air was ambient air, which was catalytically cleaned at 400°C. The flow through the sampler was controlled by the high vacuum of the PTR detector, which resulted in a sampler outlet flow of about 100 ml/min.

### Turbulent flow mixing elements

Aerodynamic mixing elements were added in the flow path of the sampler inside the first chamber for real-time analysis to obtain turbulent flow for better gas exchange and optimal mixing of the sweat volatile headspace volume.

### Controlled temperatures

The sampler was kept under the armpit and achieved body temperature after a short conditioning time. The sampling tubing from the outlet of the sampler to the inlet of the PTR detector was heated by a heating wire, insulation materials, and insulation tape. This setup allowed for the required flexibility for a person moving inside the experiment area and allowed for the sampling tubes to be intertwined multiple times. The sampling line tubing was heated to 55°C to minimize carry-over and washout times between experiments. It was very important to avoid cold spots, which could result in significant artefact spikes through aerosol release.

### Cleaning of the sampler

It was important that all of the materials in contact of the flow path from the source to the detector were readily cleanable. The sampler and custom Teflon connector were cleaned in a methanol bath with ultrasonics for 20 minutes before each experiment. The heated PFA lines were flushed out with clean gas for 30 minutes before the experiment.

### Sampler comfort

The sampler could be carried sufficiently comfortably on the body for prolonged sampling periods (i.e. up to 60 minutes). A very simple version of an armpit sampler for real-time monitoring of volatile organic compounds has been described previously [66].

### Synchronous Acquisition

While real-time mass spectrometry is uniquely powerful for observing dynamic processes, it lacks structural identification capability and cannot distinguish between isomeric compounds. To address this limitation, we collected sweat volatiles from the same skin area using time-integrative HiSorb samplers. These samples were thermally desorbed into a comprehensive two-dimensional gas chromatograph, an effective technique for separating volatile chemicals from complex mixtures and for resolving isomeric compounds.

### Artefact Detector

An internal standard mixture containing 13 VOCs was pre-mixed within the PTR-QTOF instrument using a constant flow of clean air generated by a catalytic converter maintained at 400 °C. The VOC mixture was obtained from Apel-Riemer Environmental in a certified gas cylinder at a nominal concentration of 1 ppmv dispersed in nitrogen. During the experiment, the gas was introduced at a mixing ratio of 1 mL min^-1^ of standards into 100 mL min^-1^ of clean air. The selected gas mixtures contained the following VOCs standards: methanol, acetonitrile, acetaldehyde, acrylonitrile, acetone, isoprene, 2-butanone, benzene, toluene, m-xylene, 1,2,4-trimethylbenzene, alpha-pinene, and beta-caryophyllene were used at final concentrations of 10 ppbv (except beta-caryophyllene, which was at 1 ppbv). This step proved to be essential for good quality control across the sampling runs with regard to possible leaks and washout behaviors related to sampler surface interactions.

### Real-time VOC Analysis (PTR-QTOF)

#### Instrumentation

The chemical concentrations emitted from the skin and the sweat glands were measured in real-time with a proton-transfer-reaction time-of-flight high-resolution mass spectrometer (PTR-QTOF, Vocus PTR [67], TOF Werk, Thun, Switzerland), model Vocus 2R. This model has a mass resolution of approximately 14000, which is sufficient to distinguish the vast majority of isobaric VOCs occurring in nature. The reactant we used was ion-exchanged water (MilliQ). This instrument has been predominantly employed to monitor environmental air samples; however, we have previously demonstrated its utility in human subjects within the context of breath analysis [54,68].

### Offline VOC Collection and Analysis (HiSorb-GC×GC-QTOF)

#### HiSorb sampling probes

Volatile compounds were collected using HiSorb™ probes (Markes International, Llantrisant, UK). Each stainless-steel probe (75 mm length) was coated with a polydimethylsiloxane/divinylbenzene (PDMS/DVB) copolymer film, providing a total sorptive volume of 65 µL [69]. Probes were conditioned before use in a TC-20 tube conditioner at 250 °C for 1 h under a nitrogen flow of 100 mL min^-1^, then stored in stainless-steel desorption tubes with mesh inserts and sealed using Swagelok™ caps. For sampling, probes were handled with thermally treated tweezers (150 °C, 20 min) and mounted on screw-on holders. A custom 3D-printed polycarbonate sampler housed both the HiSorb™ probe and an adjacent chamber for real-time sweat VOC monitoring. Probes were exposed for 10 min to collect sweat VOCs, while blanks were exposed for the same duration with the sampler sealed by a plastic lid. After exposure, probes were returned to desorption tubes and stored at −20 °C until analysis. All samples were analyzed within 12 h of collection to ensure sample integrity.

#### GC×GC-QTOF analysis

Thermal desorption of the HiSorb probes was performed on a Markes Centri multi-mode platform using a two-step process. A cold trap (C4–C32 range) positioned between the desorption tube and the chromatographic column concentrated analytes. Pre-purging at 50 mL min^-1^ for 2 min minimized humidity effects. Primary desorption was conducted at 35 mL min^-1^, with temperature ramped from 60 °C to 300 °C and held for 5 min. The cold trap was maintained at 2 °C, then flash-heated to 300 °C for 7 min to release analytes. Trap desorption employed a split flow of 3.5 mL min^-1^ and a column flow of 0.5 mL min^-1^; the split fraction was collected for subsequent analysis. Post-run heating at 300 °C for 5 min minimized carryover. Comprehensive GC×GC-QTOF was carried out on an Agilent 7890B system equipped with DB-5MS and DB-INNOWAX (Agilent) columns operated in constant-flow mode. The first column was set to 0.5 mL min^-1^ and the second to 18 mL min^-1^, with a 2 s modulation period. The modulator volume (19 µL) yielded 5-7 sub-peaks per 1D feature, facilitating Gaussian fitting to reconstruct first-dimension retention times (tR1). A passive splitter directed the second-dimension effluent to an Agilent 7250 QTOF and an inactive FID at a 6:1 ratio, ensuring compatibility with the QTOF’s flow capacity of <3 mL min^-1^. Re-collected samples were used for the compound identification. Ten sweat samples and an equal number of blanks from the sampling campaign were pooled for comparability; due to unequal sample availability, the final dataset comprised eight pooled sweat and three pooled blank samples (n = 11).

### Physiological Data Acquisition

We monitored the participants’ sympathetic nervous system (SNS) response during the experimental sessions by recording the electrodermal activity (EDA), a commonly used source to indirectly infer the sympathetic activations of the autonomic nervous system by means of non-invasive peripheral measurements [48]. Specifically, the EDA is a measure of variations over time in the electrical impedance of the skin due to the sweat gland activity, mediated by the sympathetic nervous system (SNS) [49]. We acquired the participants’ EDA signals using two standard Ag/AgCl electrodes placed on the third phalanx of the middle and index fingers of the non-dominant hand. We used Shimmer3 GSR+ unit^1^ wearable devices to acquire the EDA at a sampling frequency of 250 Hz. To mitigate the presence of artifacts or spurious skin conductance variations, we asked participants to keep the hand equipped with the wearable device as steady as possible and to refrain from speaking throughout the emotional stimulation sessions.

## Data Preprocessing and Feature Extraction

### Physiological Data Modelling and Fear Index Estimation

#### EDA Preprocessing

To estimate the Fear Index (FI), we used the physiological and behavioural data pertaining only to the fear-inducing session of the VR scenario. Therefore, we first segmented the EDA signals by retaining only the temporal window corresponding to the fear-inducing session. The EDA signal contains information on two different components [49]: the skin conductance level (SCL) and the skin conductance response (SCR). The SCL is the slow-varying baseline that reflects the general psychophysiological state of the person. On the contrary, the SCR consists of the superimposed high-frequency variations of the signal arising in response to external stimuli or as spontaneous activations. Each SCR originates from a neural burst in the sudomotor nerve activity (SMNA), the neural driver of the short-term electrodermal response to the stimulus. To derive these components from the raw EDA signals, we first low-pass filtered (2 Hz) the signals with a zero-phase forward and reverse eight-order digital IIR Butterworth filter to suppress noise and movement artefacts. Subsequently, we downsampled signals to 50 Hz and normalized them via z-scoring to facilitate the convergence of subsequent processing steps. Then, we applied the cvxEDA decomposition algorithm [70] to the normalized EDA signals to estimate the SCL, SCR, and SMNA time courses.

#### Feature Extraction

To model the subjective response to discrete stimuli at different levels of induced fear using the stimulus-evoked SNS response, we processed the SCL and SMNA signals to extract physiological correlates of the sympathetic branch of the nervous system corresponding to the onset of discrete stimuli. Specifically, based on the previously established psycho-physiological model of subjective fear described in [23], we computed two features: the standard deviation of the SCL signal (SCLstd) and the sum of the amplitudes of the SMNA (SMNAampsum). According to [23], we used 20-s windows for the SCL-based feature, due to its low-frequency content, and 5-s windows for the SMNA feature, due to its faster temporal dynamics. For each feature, the temporal window started at the onset of the discrete stimuli the participant encountered in the fear-inducing session, which were automatically mapped by the VR scenario.

#### LMER Model

Following the same methodological approach we developed in Baldini et al. [23], we modeled the relationship between the self-assessment of subjective fear (SASF) of the discrete events encountered in the VR scenario—rated afterwards by participants—as a function of the SNS dynamics described by the EDA features. This allowed us to take advantage of the previously validated psycho-physiological model [23] used to map the relationship between subjective fear and EDA dynamics and to fit it to the newly acquired data pertaining to this study. We purposely adopted a mixed-effects regression to account for the hierarchical structure of the experimental protocol, given by the repeated measurement design spanning both participants and discrete stimuli. Indeed, the mixed-effects design enabled us to model the relationship between physiological features and the fear score, accounting for the variability within and across participants and stimulus contents. Thus, we explored the general linear relationships between the SASF and physiological features without violating the independence assumptions of linear regression.

More in detail, we performed a linear mixed-effects regression (LMER) analysis where the SASF was the dependent variable and the physiological features (i.e., SCLstd and SMNAampsum) were the interacting fixed-effects, with the participant and discrete stimuli identifiers modelled as independent random intercept effects. Therefore, we adopted the following model:

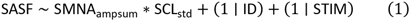

with ID and STIM corresponding to the participant and discrete stimulus identifiers, respectively. By fitting the model in Eq. 1, we obtained the estimates of three coefficients, i.e., β1, β2, and β3, to map the linear relationship between the SASF and SMNAampsum, SCLstd, and the interactions among SMNAampsum and SCLstd, respectively. All analyses concerning the LMER models were executed using the R package lme4 [71].

#### Continuous Fear Index

Using the LMER fit results, we devised a continuous estimate of the FI. Firstly, we computed the SMNAampsum and SCLstd measures over time in non-overlapping temporal windows of 5 s. Specifically, the measures were computed on segments of 700 s spanning the temporal range of 100 s before the emotional stimulation started until its end. Then, based on the fixed-effects coefficients of the LMER fit, we computed the continuous FI variable according to the following equation:

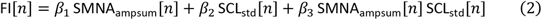

with n being the temporal index. This corresponds to the most reliable estimate of an objective time-resolved physiology-informed fear index based on a previously established relationship [23] between sympathetic dynamics and the behavioural response to fearful events.

### Real-time Chemical Data Preprocessing

#### Chemical Preprocessing

Data processing of the chemical signals from the PTR-QTOF was performed using Tofware 3.2.5 (TofWerk, Thun, Switzerland). We applied a non-targeted analysis workflow to each sample, which involved several key steps. First, a background spectrum was defined for baseline correction, noise reduction, and control of instrumental drift. This was followed by peak-shape refinement to eliminate artefacts and to prepare the data for high-resolution fitting to the time-to-mass function. Subsequently, high-resolution fitting was performed through recalibration of the m/z axis to enable accurate peak separation across the mass spectrum. An automated peak-finding algorithm was then used for peak detection to identify relevant signals. Finally, the individual sample tables were compiled into a single data table, incorporating all detected features across all participants with an intensity above a threshold of at least 10 ions per second.

#### Elemental formula assignments

Elemental compositions were determined for each accurate mass feature using the formula assignment tool in Tofware 3.2.5, which operates on a statistical probability basis. The assignment was based on a set of selected elements, primarily carbon, hydrogen, oxygen, and nitrogen, but also included less abundant elements commonly found in volatile organic compounds, such as sulfur, silicon, or halogens. The probability of each elemental composition was evaluated using a combination of four criteria. These included the assignment accuracy, which quantified the difference between the theoretical and measured m/z values, and the isotope peak ratio, which assessed the deviation between theoretical and observed isotopic peak heights. Additionally, we considered the chemical feasibility of the proposed formula and examined highly correlated features within the dataset that exhibited unambiguous mass differences—such as water adducts with a mass difference of 18.011 Da or specific, well-characterized ion fragments.

#### Data normalization

Data normalization with Tofware 3.2.5 across the whole sampling batch was not reproducible. These issues could not be resolved by the software developer Aerodyne Research, Inc. (Aerodyne Research, Inc., Billerica, MA, USA). However, the continuous addition of internal standards with our sampling system allowed us to normalize all data relative to xylene, which resolved the data normalization issue.

#### Data exclusion (mass features)

We reduced the initial set of 234 mass features to the first 183 features and finally to 129 features. A comparison of the integrated PTR-QTOF time traces for the last 30-seconds of the 100-second baseline was compared with the last 30 seconds of steady-state conditions of the empty sampler (t-tests with p.adj. < 0.01 and FC > 2.0) adjusted using a false discovery rate (FDR - Benjamini-Hochberg) correction, reduced the mass features from 234 to 183 features. Additionally, a comparison of the integrated PTR-QTOF time traces for the 10-min fear-induction phase was compared with the cumulative 5-min resting baseline and 5-min neutral stimulation phase (FC > 1.01) to reduce the mass features from 183 to the final 129 features used for further data analysis, modeling, and statistical analysis.

#### Data exclusion (participants)

Eight participants were excluded from the final joint chemical– physiological modelling dataset. Three participants were excluded because of high residual sweat signals that produced prolonged chemical washout and interfered with the fear-induction recording (S13, S20, S26), most likely due to insufficient cleaning of the axillary sampling area before the experiment. Two participants were excluded because of leaks during jump-scare events in the fear scenario (S09, S27), which produced abrupt signal drops and required several minutes for the sampling system to return to steady-state conditions. One participant was excluded because of an instrumental issue involving unheated sampling tubes (S41). Finally, two additional participants were excluded because of noisy chemical data (S04) and failure of the EDA recording system (S33), respectively. These exclusions resulted in a final dataset of 37 healthy participants for the joint PTR-QTOF/CD-ICA/HBEN analysis (14 males, 23.4 ± 2.3 years; 23 females, 23.9 ± 2.7 years).

#### Nuisance Variable Regression

We used a General Linear Model (GLM) to regress out the variance associated with known confounding factors (nuisance variables). This was performed independently for each chemical time series. For each *Y_i_*, a GLM was fitted using three robust calibrant time-series (*X*) as predictors, with the following equation (3):

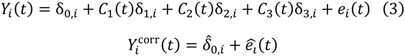

where *δ*_0,*i*_, *δ*_1,*i*_, *δ*_2,*i*_, *δ*_3,*i*_ are the regression coefficients, and *e_i_*(*t*) represents the residual error. The calibrant-corrected time series was reconstructed as the sum of the model’s intercept and the residual time series. This procedure effectively removes the variance *Y_i_* that is linearly dependent on the three calibrant time-series.

## Statistical Analysis: Identification of the Fear Fingerprint

### Chemical-Domain Group Independent Component Analysis (CD-ICA)

#### Rationale

Multi-subject ICA methods have been extensively developed, primarily in fMRI research [19]. In this scenario, the goal is to estimate functionally connected networks by decomposing linearly mixed fMRI signals into spatially independent patterns, providing both subject-specific spatial maps (SMs) and time courses (TCs). Here, we adapted an extensively validated, robust, and accurate method used in fMRI analysis to estimate subject-specific chemical TCs and SMs, based on the following assumptions:

1. The chemical composition of independent components is shared across participants.
2. Each subject has their own time courses for the chemical components, considering that the exploration of the fear scenario varied across individuals.

#### Key Steps

Among the several methods proposed in [19], we specifically employed the GICA3 back-reconstruction method, with subject-specific PCA and noise-free ICA. This specific approach provides the most robust and accurate estimates of SMs and TCs, in addition to offering an intuitive interpretation of components [19]. The procedure consisted of the following steps (see Fig. 2a):

1. Each chemical time series was scaled to have a zero mean.
2. For each subject, the time dimension (T) was reduced to T1 using subject-specific PCA. This step preserves subject-specific variance by projecting the final group solution into unique subject column spaces.
3. Subject-specific PCA-reduced data were temporally concatenated and further reduced from MT1 to T2 using group-level PCA to find a common column space.
4. Group ICA was performed on the temporally reduced group data to obtain independent group spatial maps. We estimated 40 components.
5. The GICA3 back-projection method was applied to estimate the final subject-specific TCs and SMs.

This multi-stage procedure ensures that the two key assumptions (shared chemical composition and unique subject time-courses) are effectively addressed.

The extracted components were expected to comprise both biologically meaningful independent chemical signatures and non-physiological sources of variability, including sensor-or subject-related motion, motion/leak transients, instrumental noise or fluctuations, and chemically uninterpretable patterns. Therefore, before statistical modelling, all components were subjected to a blind quality-control screening. Components showing clear artefactual characteristics or lacking a coherent chemical structure consistent with interpretable ion/adduct patterns were excluded from subsequent analyses. The remaining 29 components were retained for downstream analysis.

#### Hierarchical Bayesian Elastic Net Model (HBEN)

To identify the predictors of fear index from the set of pre-computed Independent Components (ICs), i.e., the potential fear chemosignals, we developed a custom hierarchical Bayesian regression model. The architecture of this model was specifically designed to address the key statistical challenges inherent in our high-dimensional, longitudinal dataset:

- **Hierarchical Data Structure:** To account for the non-independence of observations arising from multiple measurements being nested within each subject, we employed a hierarchical (or mixed-effects) structure with subject-specific intercepts. This allows the model to capture inter-subject heterogeneity in baseline responses while enabling population-level inferences.
- **Temporal Autocorrelation:** To handle the serial dependence between consecutive measurements within a subject’s time series, we explicitly modeled the residuals using a first-order autoregressive (AR(1)) process. This step is critical for obtaining unbiased estimates of parameter uncertainty by ensuring that the model’s innovations (i.e., the corrected residuals) satisfy the assumption of independence.
- **High-Dimensionality and Multicollinearity:** To perform robust variable selection in a setting where the number of predictors (p = 29) is substantial, and these predictors may be intercorrelated, we implemented Elastic Net regularization within the Bayesian framework. This approach combines an L1 penalty to induce sparsity (by shrinking irrelevant coefficients towards zero) and an L2 penalty to manage multicollinearity. The L2 component confers a “grouping effect,” which encourages the model to treat correlated predictors as a group, leading to more stable and interpretable variable selection than the L1 penalty alone.

The entire model was specified within a probabilistic programming framework, allowing for full Bayesian inference. This provides complete posterior distributions for every parameter, offering a comprehensive quantification of uncertainty for our conclusions.

#### Probabilistic Model Specification

The model is defined as a generative hierarchy of probability distributions. Both the predictor variables (ICs) and the response variable were standardized (Z-scored) prior to model fitting to a common scale, facilitating the specification of meaningful priors.

#### Likelihood Function

The generative process for an observation *y_t_* at time *t* for subject *s*(*t*) is decomposed into a deterministic mean component, *μ_t_*, and a serially correlated error term, *ν_t_*. This is formally expressed as:

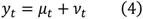

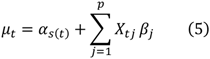

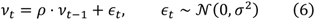

Where p=29 is the number of CD-ICA component time courses entered as predictors, *α_s_*_(*t*)_ is the subject-specific intercept, *X_t_*_j_ is the value of the j-th predictor, *β*_j_ is its associated coefficient, *ρ* is the first-order autocorrelation parameter, and *ε_t_* are the independent and identically distributed innovations with standard deviation *σ*. By substituting these definitions, we can formulate the complete likelihood for the model. The formulation explicitly handles the initialization of the AR(1) process for the first observation of each subject to ensure stationarity:

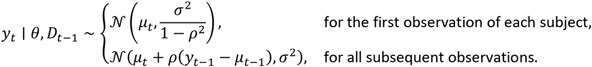

where *θ* represents the full set of model parameters and *D_t_*_–1_ represents the history of observations up to time *t* − 1.

#### Prior Distributions

We adopted a strategy of using weakly informative priors, which provide gentle regularization to prevent inferentially unstable or unrealistic parameter estimates, while still allowing the data to dominate the final posterior estimates.

#### Subject Intercepts (*α_s_*)

The subject-specific intercepts were modeled as draws from a common population distribution, which is the cornerstone of the hierarchical approach. A standard Normal prior was used, consistent with the standardized scale of the response variable:

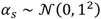

#### Autocorrelation Coefficient (*ρ*)

The prior for *ρ* was a Normal distribution centered at zero, which does not presuppose positive or negative autocorrelation, with a standard deviation chosen to assign most of the prior mass within the valid range of [−1,1]:

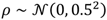

#### Innovation Variance (*σ*)

A Half-Normal prior was assigned to the standard deviation of the innovations, constraining it to be positive and weakly favoring smaller values, which is a reasonable assumption for the residual variance of a predictive model on standardized data:

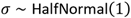

#### Regression Coefficients (*β*_j_) and Elastic Net Regularization

We implemented Elastic Net regularization via a modular, two-stage prior structure. First, a deliberately wide, weakly informative Normal prior was assigned to each coefficient as a baseline:

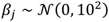

This wide prior ensures that the posterior estimates are not overly constrained by the prior itself; a coefficient of 10 on standardized data would represent an implausibly large effect. The regularization is then introduced by augmenting the model’s log-posterior probability directly with two penalty terms, controlled by hyperparameters *λ_L_*_1_ and *λ_L_*_2_.

#### Regularization Hyperparameters (*λ_L_*_1_, *λ_L_*_2_)

Instead of fixing the penalty strengths a priori, we inferred them from the data by assigning them their own hyperpriors. An Exponential distribution was chosen as it places more prior weight on smaller penalty values (less regularization) while having a long tail that allows for strong regularization if supported by the data:

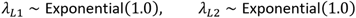

This fully Bayesian approach allows the model to learn the optimal balance between sparsity and grouping directly from the data structure.

#### Full Posterior Distribution

Combining these components via Bayes’ theorem, the joint posterior distribution for the complete set of parameters *θ* = {β, α, *ρ*, *σ*, *λ_L_*_1_, *λ_L_*_2_} is proportional to the product of the likelihood and all prior distributions, further shaped by the Elastic Net penalty potentials:

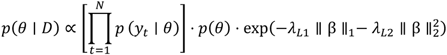

where *p*(*θ*) is the product of all parameter priors, and ∥ β ∥_1_ and ∥ β ∥^2^_2_ are the L1 and squared L2 norms of the coefficient vector, respectively.

#### Posterior Inference and Implementation

The joint posterior distribution is analytically intractable. Therefore, we performed inference using Markov Chain Monte Carlo (MCMC) sampling. The model was implemented in the PyMC probabilistic programming language (v5.0 or later). We used the No-U-Turn Sampler (NUTS), a highly efficient gradient-based Hamiltonian Monte Carlo algorithm, to draw samples from the posterior distribution. We ran four independent MCMC chains in parallel, with each chain generating 4,000 total iterations. The first 2,000 iterations of each chain were discarded as a warm-up (tuning) phase, and the subsequent 2,000 were retained for inference, yielding a total of 8,000 effective samples from the joint posterior.

Convergence of the MCMC chains to the stationary posterior distribution was rigorously assessed. We confirmed that the Gelman-Rubin diagnostic statistic (R^) was less than 1.01 for all model parameters, indicating successful convergence.

#### Criterion for Predictor Relevance

A predictor IC was identified as “credibly relevant” if its corresponding coefficient *β*_j_ was statistically distinguishable from zero. For this, we adopted a Bayesian decision rule: the 95% Highest Density Interval (HDI) of the marginal posterior distribution for *β*_j_ must not contain zero. This criterion provides strong probabilistic evidence that the predictor exerts a non-zero effect (either strictly positive or strictly negative) on the response variable.

### GC×GC Data Analysis

#### Preprocessing

Raw data files were processed to generate images containing all chromatographic and spectral information using GC Image (version 2.8). Images were automatically preprocessed by applying a spectral filter to remove background ions, followed by automatic phase correction and peak detection. A cumulative image was then created for compound identification by superimposing all fear samples from the subjects, while individual images were later used to selectively extract responses using extracted ion images. The full analytical approach and workflow are described in detail in our previous publication [35].

#### Compound identification

We identified the m/z features detected with GC×GC-QTOF based on our previously published approach [35]. Chemical identification is based on a combination of spectral library searches, linear retention index matching, and assignment of elemental formulae to the molecular or characteristic ions, possibly due to the high mass resolution power of the mass spectrometer.

## Statistical Analysis

Following data extraction from the individual samples, a data matrix containing 296 chemical features across 116 samples was generated. Data were first normalized to the internal standard and subsequently filtered through pairwise comparisons to remove compounds present in blank samples. Filtering was performed using volcano plot analysis (p-value significance versus fold change), with thresholds of fold change > 3 and significance level p < 0.01, adjusted using a false discovery rate (FDR - Benjamini-Hochberg) correction. This procedure yielded a final list of 166 identified compounds.

## Data and code availability

The chemical, physiological, and psychometric data, the analysis code (Python and MATLAB), and all materials will be made available on a public repository (<u>OSF</u>). The CD-ICA pipeline, the hierarchical Bayesian Elastic-Net model, and the robustness-analysis scripts will be deposited as stand-alone, annotated codebases with instructions for reproduction.

## Author contributions

T.B., A.G., E.P.S. and F.D.F. conceived the study. T.B., A.G., M.R., N.V. and A.L.C. designed the experimental and analytical strategy. T.B. developed the axillary sampling and real-time chemical measurement approach, with support from M.R., F.M.V., D.B. and T.L. A.G. developed the statistical and physiological modelling strategy, with support from A.L.C., A.Ga. T.B., M.R., A.L.C., A.Ga., S.R. and M.P. performed the experiments. T.B., A.G., M.R., A.L.C., A.Ga. and S.R. analysed the data. T.B., M.R., F.M.V., T.L. and D.B. contributed to the development and implementation of the instrumental methodology. A.G., A.L.C. and A.Ga contributed to the development and implementation of the data-analysis methodology. T.B. and A.G. wrote the original draft. All authors reviewed and edited the manuscript. A.G., F.D.F. and E.P.S. contributed to funding acquisition. T.B., A.G., E.P.S. and F.D.F. contributed to project administration. E.P.S. and F.D.F. supervised the work. All authors read and approved the final manuscript.

## Supporting information

Supplementary Informations

## Acknowledgements

We thank Ilona Croy, Thomas Hummel and Sean Froese for their thoughtful and constructive feedback on an earlier version of this manuscript, which helped us refine the interpretation of the findings and clarify the scope of the mechanistic claims. We thank Veronika Pospisilova and the team at TOF Werk for testing our early version of the sampler at TOF Werk in Thun, Switzerland. We thank Nuno Gomes and Gün R. Semin for early discussions on participant preparation for body-odour sampling. We acknowledge CISUP, University of Pisa, for supporting the acquisition of the Vocus CI-TOF instrument used in this work. We also thank the members of the POTION consortium for discussions that helped shape the experimental framework. This work was supported by the European Union Horizon 2020 research and innovation programme through the POTION project.

## Footnotes

1 https://www.shimmersensing.com/product/shimmer3-gsr-unit/

