## Supplementary Informations for "A time-resolved axillary volatile fingerprint of human fear"

**Contents**

**Supplementary Table 1 |** Detected volatile organic compounds

**Supplementary Table 2 |** Compound identification of fear-associated CD-ICA components

**Supplementary Table 3 |** Compound properties

**Supplementary Table 4 |** Subjective assessment of the VR fear-induction scenario

**Supplementary Table 5 |** Physiological assessment of the VR fear-induction scenario

**Supplementary Fig. 1 |** STAI-Y1 scores after neutral and fear sessions

**Supplementary Fig. 2 |** Self-reported fear scores after neutral and fear sessions

**Supplementary Fig. 3 |** EDASymp during neutral and fear sessions

**Supplementary References**

Unless otherwise specified, chemical analyses refer to the final axillary volatilome dataset after exclusion criteria (37 participants). Manipulation checks were computed on all available enrolled participants: subjective ratings were available for 45 participants, whereas the electrodermal sympathetic activity was measurable for 44 participants because of one EDA recording failure. The hierarchical Bayesian fear-index model was fitted on the subset of participants with valid chemical and EDA-derived fear-index time series.

### Supplementary Table 1: Detected volatile organic compounds

**Supplementary Table 1 | Detected volatile organic compounds (*m/z* features with PTR-QTOF) for the 45 healthy participants (28 females, mean age 23.5 ± 2.4 years; 17 males, mean age 23.7 ± 2.2 years).**

| Nr. | m/z value | Ion formula | Baseline vs. blank | | Fear vs. baseline | Inclusion |
|---|---|---|---|---|---|---|
| | | | FC > 2.0 | p.adj < 0.01 | FC > 1.01 | |
| 1 | 29.997 | NO+ | | | | no |
| 2 | 31.018 | CH3O+ | | | | no |
| 3 | 31.989 | O2+ | | | | no |
| 4 | 33.033 | CH5O+ | | | | no |
| 5 | 34.029 | H4NO+ | | | | no |
| 6 | 37.028 | H5O2+ | | | | no |
| 7 | 38.015 | C3H2+ | 3.03 | 7.51E-15 | | no |
| 8 | 39.023 | C3H3+ | 3.61 | 6.44E-15 | 1.02 | yes |
| 9 | 41.039 | C3H5+ | 3.63 | 7.12E-14 | 1.02 | yes |
| 10 | 42.034 | C2H4N+ | 7.00 | 1.37E-26 | | no |
| 11 | 43.018 | C2H3O+ | 3.05 | 1.15E-13 | 0.97 | yes |
| 12 | 43.054 | C3H7+ | 2.42 | 6.18E-14 | 1.03 | yes |
| 13 | 44.049 | C2H6N+ | 2.95 | 7.95E-19 | 1.04 | yes |
| 14 | 45.033 | C2H5O+ | 2.81 | 2.33E-17 | | no |
| 15 | 45.992 | NO2+ | | | | no |
| 16 | 46.029 | CH4NO+ | | | | no |
| 17 | 46.065 | C2H8N+ | 4.61 | 3.59E-14 | 1.12 | yes |
| 18 | 47.013 | CH3O2+ | 2.17 | 7.61E-12 | 0.99 | yes |
| 19 | 47.049 | C2H7O+ | 3.21 | 3.24E-15 | 0.92 | yes |
| 20 | 48.008 | H2NO2+ | | | | no |
| 21 | 50.015 | C4H2+ | 4.27 | 1.05E-24 | 0.98 | yes |
| 22 | 51.044 | CH7O2+ | 3.44 | 5.15E-16 | 0.97 | yes |
| 23 | 53.003 | Unknown | 4.36 | 5.39E-17 | | no |
| 24 | 53.039 | C4H5+ | 4.70 | 8.88E-19 | 0.96 | yes |
| 25 | 54.034 | C3H4N+ | 11.55 | 1.24E-23 | | no |
| 26 | 55.039 | H7O3+ | 2.48 | 1.06E-11 | 1.04 | yes |
| 27 | 57.033 | C3H5O+ | 4.14 | 3.42E-15 | 1.02 | yes |
| 28 | 57.070 | C4H9+ | 6.53 | 2.75E-14 | 0.97 | yes |
| 29 | 59.049 | C3H7O+ | 10.63 | 1.16E-19 | 1.18 | yes |
| 30 | 60.021 | C2H4O2+ | 3.43 | 7.83E-19 | 1.07 | yes |
| 31 | 61.028 | C2H5O2+ | 4.89 | 2.45E-15 | 0.96 | yes |
| 32 | 63.044 | C2H7O2+ | 3.34 | 1.40E-15 | | no |
| 33 | 64.003 | H2NO3+ | | | | no |
| 34 | 65.039 | C5H5+ | 5.24 | 7.06E-21 | 0.98 | yes |

| | | | | | | |
|---|---|---|---|---|---|---|
| 35 | 65.060 | C2H9O2+ | 3.72 | 1.75E-13 | 0.95 | yes |
| 36 | 66.046 | C5H6+ | 4.58 | 1.11E-25 | 0.97 | yes |
| 37 | 67.054 | C5H7+ | 5.17 | 2.11E-16 | 0.97 | yes |
| 38 | 68.026 | C4H4O+ | 4.21 | 3.62E-28 | | no |
| 39 | 68.062 | C5H8+ | 4.85 | 9.24E-28 | 0.98 | yes |
| 40 | 69.033 | C4H5O+ | 2.04 | | 1.02 | yes |
| 41 | 69.055 | CH9O3+ | 4.55 | 4.07E-13 | 0.97 | yes |
| 42 | 69.070 | C5H9+ | 4.72 | 6.59E-22 | | no |
| 43 | 71.013 | C3H3O2+ | 3.32 | 2.62E-12 | | no |
| 44 | 71.049 | C4H7O+ | 5.07 | 1.20E-16 | | no |
| 45 | 71.086 | C5H11+ | 7.74 | 1.07E-15 | 0.97 | yes |
| 46 | 72.044 | C3H6NO+ | 10.04 | 1.85E-20 | | no |
| 47 | 73.028 | C3H5O2+ | 2.81 | 4.07E-12 | | no |
| 48 | 73.050 | H9O4+ | 2.45 | 8.26E-08 | 1.05 | yes |
| 49 | 73.065 | C4H9O+ | 5.85 | 1.06E-27 | | no |
| 50 | 74.024 | C2H4NO2+ | 2.60 | 1.15E-11 | | no |
| 51 | 74.071 | C2H8N3+ | 4.78 | 1.37E-26 | | no |
| 52 | 75.044 | C3H7O2+ | 5.09 | 9.21E-12 | | no |
| 53 | 76.076 | C3H10NO+ | 3.23 | 5.91E-13 | 1.13 | yes |
| 54 | 77.060 | C3H9O2+ | 9.71 | 3.16E-15 | 1.15 | yes |
| 55 | 78.046 | C6H6+ | 7.08 | 5.42E-24 | | no |
| 56 | 79.039 | C2H7O3+ | 8.87 | 2.67E-09 | 0.95 | yes |
| 57 | 79.054 | C6H7+ | | | | no |
| 58 | 81.045 | C4H5N2+ | 0.27 | | 1.02 | yes |
| 59 | 81.070 | C6H9+ | 10.66 | 3.15E-21 | 0.96 | yes |
| 60 | 83.049 | C5H7O+ | 3.31 | 1.86E-11 | | no |
| 61 | 83.086 | C6H11+ | 8.84 | 1.90E-15 | 0.94 | yes |
| 62 | 84.021 | C4H4O2+ | | | | no |
| 63 | 84.044 | C4H6NO+ | 2.64 | 2.10E-12 | | no |
| 64 | 84.057 | C5H8O+ | | | | no |
| 65 | 85.028 | C4H5O2+ | 4.16 | 1.04E-12 | 1.01 | yes |
| 66 | 85.065 | C5H9O+ | 3.62 | 1.15E-13 | | no |
| 67 | 85.101 | C6H13+ | 8.70 | 3.88E-17 | 0.96 | yes |
| 68 | 86.036 | C4H6O2+ | 2.94 | 3.89E-19 | | no |
| 69 | 86.060 | C4H8NO+ | 5.17 | 4.32E-22 | 0.98 | yes |
| 70 | 87.044 | C4H7O2+ | 3.92 | 4.12E-12 | | no |
| 71 | 87.080 | C5H11O+ | 2.58 | 2.86E-09 | 1.20 | yes |
| 72 | 88.039 | C3H6NO2+ | 3.56 | 5.85E-13 | 1.04 | yes |
| 73 | 88.076 | C4H10NO+ | 3.32 | 6.92E-15 | 1.12 | yes |
| 74 | 89.023 | C3H5O3+ | 3.93 | 7.85E-12 | | no |
| 75 | 89.060 | C4H9O2+ | 8.14 | 3.99E-17 | 0.98 | yes |
| 76 | 90.091 | C4H12NO+ | 3.87 | 2.95E-23 | | no |
| 77 | 91.042 | C6H5N+ | 3.82 | 8.94E-06 | 1.13 | yes |
| 78 | 91.054 | C7H7+ | 6.20 | 6.83E-26 | 0.98 | yes |
| 79 | 91.075 | C4H11O2+ | 6.18 | 9.01E-22 | 0.98 | yes |
| 80 | 92.062 | C7H8+ | 6.46 | 6.03E-28 | 0.97 | yes |
| 81 | 93.057 | C6H7N+ | 5.10 | 5.27E-05 | 0.97 | yes |
| 82 | 93.070 | C7H9+ | 7.08 | 6.03E-28 | 0.99 | yes |
| 83 | 95.049 | C6H7O+ | 6.79 | 6.99E-14 | | no |
| 84 | 95.070 | C3H11O3+ | 0.14 | | 1.11 | yes |
| 85 | 95.086 | C7H11+ | 17.87 | 5.39E-17 | 0.97 | yes |
| 86 | 97.028 | C5H5O2+ | 3.93 | 1.66E-10 | 1.01 | yes |
| 87 | 97.050 | C2H9O4+ | 4.21 | 2.33E-08 | 0.93 | yes |
| 88 | 97.065 | C6H9O+ | | | | no |
| 89 | 97.101 | C7H13+ | 3.29 | 7.03E-10 | 0.94 | yes |

| | | | | | | |
|---|---|---|---|---|---|---|
| 90 | 99.044 | C5H7O2+ | 3.56 | 1.44E-11 | 1.01 | yes |
| 91 | 99.080 | C6H11O+ | 3.16 | 8.64E-12 | | no |
| 92 | 99.117 | C7H15+ | 2.07 | | | no |
| 93 | 100.039 | C4H6NO2+ | 3.25 | 2.23E-13 | 1.02 | yes |
| 94 | 101.023 | C4H5O3+ | 4.39 | 5.24E-14 | 1.01 | yes |
| 95 | 101.060 | C5H9O2+ | 3.78 | 1.19E-11 | | no |
| 96 | 101.096 | C6H13O+ | | | | no |
| 97 | 102.055 | C4H8NO2+ | 3.84 | 4.41E-17 | 1.02 | yes |
| 98 | 103.039 | C4H7O3+ | 5.21 | 2.87E-10 | | no |
| 99 | 103.075 | C5H11O2+ | 2.01 | 1.65E-13 | | no |
| 100 | 105.018 | C3H5O4+ | 3.33 | 4.75E-10 | 1.02 | yes |
| 101 | 105.055 | C4H9O3+ | 5.95 | 1.68E-07 | 1.05 | yes |
| 102 | 105.070 | C8H9+ | 4.31 | 2.05E-18 | 0.97 | yes |
| 103 | 105.091 | C5H13O2+ | | | | no |
| 104 | 106.078 | C8H10+ | 6.58 | 3.23E-28 | 0.98 | yes |
| 105 | 107.034 | C3H7O4+ | 2.93 | 1.62E-08 | | no |
| 106 | 107.070 | C4H11O3+ | 0.11 | 1.18E-05 | 1.02 | yes |
| 107 | 107.086 | C8H11+ | 12.95 | 1.55E-20 | | no |
| 108 | 109.028 | C6H5O2+ | 6.32 | 2.38E-09 | 1.03 | yes |
| 109 | 109.050 | C3H9O4+ | | | | no |
| 110 | 109.065 | C7H9O+ | 3.02 | 2.52E-08 | 0.98 | yes |
| 111 | 109.101 | C8H13+ | 9.53 | 1.09E-18 | 0.93 | yes |
| 112 | 111.044 | C6H7O2+ | 3.98 | 1.18E-11 | | no |
| 113 | 111.080 | C7H11O+ | | | | no |
| 114 | 111.117 | C8H15+ | 7.22 | 1.02E-16 | 1.03 | yes |
| 115 | 113.023 | C5H5O3+ | 11.56 | 4.46E-19 | 1.08 | yes |
| 116 | 113.060 | C6H9O2+ | 3.75 | 1.02E-12 | | no |
| 117 | 113.096 | C7H13O+ | | | | no |
| 118 | 113.132 | C8H17+ | 5.07 | 1.56E-12 | 1.03 | yes |
| 119 | 115.039 | C5H7O3+ | 2.62 | | 1.01 | yes |
| 120 | 115.075 | C6H11O2+ | 3.66 | 3.64E-09 | 1.02 | yes |
| 121 | 117.018 | C4H5O4+ | 2.40 | | | no |
| 122 | 117.055 | C5H9O3+ | 3.59 | 6.56E-11 | 1.02 | yes |
| 123 | 117.091 | C6H13O2+ | 2.93 | 6.66E-09 | | no |
| 124 | 118.053 | C7H6N2+ | 3.04 | 2.28E-13 | 1.03 | yes |
| 125 | 119.034 | C4H7O4+ | 3.35 | 2.06E-09 | | no |
| 126 | 119.070 | C5H11O3+ | | | | no |
| 127 | 119.086 | C9H11+ | 3.69 | 4.59E-11 | 0.98 | yes |
| 128 | 119.107 | C6H15O2+ | | | | no |
| 129 | 120.093 | C9H12+ | 6.26 | 2.17E-25 | 0.99 | yes |
| 130 | 121.028 | C7H5O2+ | 3.96 | | 1.06 | yes |
| 131 | 121.050 | C4H9O4+ | | | | no |
| 132 | 121.101 | C9H13+ | 12.92 | 5.25E-19 | | no |
| 133 | 123.029 | C3H7O5+ | 4.46 | 1.08E-08 | | no |
| 134 | 123.044 | C7H7O2+ | | | | no |
| 135 | 123.065 | C4H11O4+ | | | | no |
| 136 | 123.080 | C8H11O+ | 3.45 | 1.23E-20 | 0.98 | yes |
| 137 | 123.117 | C9H15+ | 8.15 | 6.29E-17 | 0.96 | yes |
| 138 | 125.023 | C6H5O3+ | 5.60 | 6.79E-11 | 1.01 | yes |
| 139 | 125.044 | C3H9O5+ | 0.28 | 4.60E-06 | | no |
| 140 | 125.060 | C7H9O2+ | 3.24 | 2.06E-12 | | no |
| 141 | 125.081 | C4H13O4+ | 0.42 | | | no |
| 142 | 125.096 | C8H13O+ | 3.79 | 5.78E-13 | 0.97 | yes |
| 143 | 125.132 | C9H17+ | 3.05 | 1.26E-13 | 0.93 | yes |
| 144 | 126.091 | C7H12NO+ | | | | no |

| | | | | | | |
|---|---|---|---|---|---|---|
| 145 | 127.039 | C6H7O3+ | 3.03 | 2.42E-08 | | no |
| 146 | 127.075 | C7H11O2+ | | | | no |
| 147 | 127.112 | C8H15O+ | 3.02 | 5.00E-10 | 0.84 | yes |
| 148 | 127.148 | C9H19+ | 2.26 | 9.56E-07 | 1.03 | yes |
| 149 | 129.018 | C5H5O4+ | | | | no |
| 150 | 129.055 | C6H9O3+ | 3.49 | 1.50E-09 | | no |
| 151 | 129.091 | C7H13O2+ | | | | no |
| 152 | 129.127 | C8H17O+ | 4.05 | 1.01E-12 | 1.08 | yes |
| 153 | 130.086 | C6H12NO2+ | | | | no |
| 154 | 131.034 | C5H7O4+ | 4.49 | 1.68E-13 | 1.03 | yes |
| 155 | 131.070 | C6H11O3+ | 3.28 | 5.28E-09 | | no |
| 156 | 131.107 | C7H15O2+ | 0.13 | | 1.02 | yes |
| 157 | 133.050 | C5H9O4+ | 3.82 | 1.34E-10 | | no |
| 158 | 133.086 | C6H13O3+ | | | | no |
| 159 | 133.122 | C7H17O2+ | | | | no |
| 160 | 135.065 | C5H11O4+ | | | | no |
| 161 | 135.080 | C9H11O+ | | | | no |
| 162 | 135.102 | C6H15O3+ | 2.35 | | 1.05 | yes |
| 163 | 135.117 | C10H15+ | 6.40 | 1.33E-07 | 0.99 | yes |
| 164 | 136.022 | C7H6NS+ | | | | no |
| 165 | 136.060 | C4H10NO4+ | 2.03 | 3.50E-09 | | no |
| 166 | 136.125 | C10H16+ | 5.26 | 3.58E-19 | 0.98 | yes |
| 167 | 137.044 | C4H9O5+ | 2.19 | 5.31E-06 | 0.99 | yes |
| 168 | 137.132 | C10H17+ | 17.80 | 1.56E-21 | 0.97 | yes |
| 169 | 139.039 | C7H7O3+ | 7.71 | 1.75E-11 | 1.06 | yes |
| 170 | 139.075 | C8H11O2+ | | | | no |
| 171 | 139.112 | C9H15O+ | 2.31 | 3.42E-09 | 0.95 | yes |
| 172 | 139.148 | C10H19+ | 6.02 | 5.97E-11 | 0.87 | yes |
| 173 | 141.018 | C6H5O4+ | 3.34 | 3.67E-12 | 1.01 | yes |
| 174 | 141.055 | C7H9O3+ | 2.26 | 5.49E-11 | 1.02 | yes |
| 175 | 141.091 | C8H13O2+ | | | | no |
| 176 | 141.127 | C9H17O+ | | | | no |
| 177 | 141.164 | C10H21+ | 2.74 | 6.83E-08 | 0.84 | yes |
| 178 | 142.053 | C9H6N2+ | | | | no |
| 179 | 143.034 | C6H7O4+ | 4.35 | 9.26E-09 | | no |
| 180 | 143.070 | C7H11O3+ | | | | no |
| 181 | 143.107 | C8H15O2+ | 3.48 | 3.06E-12 | 0.97 | yes |
| 182 | 143.143 | C9H19O+ | 2.22 | 1.75E-13 | 0.92 | yes |
| 183 | 144.068 | C9H8N2+ | | | | no |
| 184 | 145.050 | C6H9O4+ | 4.23 | 2.04E-09 | | no |
| 185 | 145.086 | C7H13O3+ | | | | no |
| 186 | 145.122 | C8H17O2+ | 3.76 | 1.45E-13 | 1.04 | yes |
| 187 | 147.029 | C5H7O5+ | 2.82 | 6.18E-11 | 1.02 | yes |
| 188 | 147.065 | C6H11O4+ | 2.91 | 8.90E-11 | 1.01 | yes |
| 189 | 147.102 | C7H15O3+ | | | | no |
| 190 | 147.117 | C11H15+ | 2.11 | | 0.95 | yes |
| 191 | 147.138 | C8H19O2+ | 3.49 | 2.31E-12 | 1.07 | yes |
| 192 | 149.044 | C5H9O5+ | 3.79 | 1.78E-12 | 1.02 | yes |
| 193 | 149.081 | C6H13O4+ | | | | no |
| 194 | 149.132 | C11H17+ | 10.51 | 3.11E-24 | | no |
| 195 | 150.112 | C6H16NO3+ | | | | no |
| 196 | 153.055 | C8H9O3+ | 3.15 | 1.41E-11 | 1.02 | yes |
| 197 | 153.091 | C9H13O2+ | 2.52 | 1.06E-09 | | no |
| 198 | 153.127 | C10H17O+ | 4.29 | 2.93E-10 | 0.97 | yes |
| 199 | 155.034 | C7H7O4+ | 2.48 | 3.17E-12 | 1.01 | yes |

| | | | | | | |
|---|---|---|---|---|---|---|
| 200 | 155.070 | C8H11O3+ | 2.09 | 1.67E-08 | | no |
| 201 | 155.107 | C9H15O2+ | 2.19 | 3.23E-06 | 0.98 | yes |
| 202 | 155.143 | C10H19O+ | 2.82 | 5.74E-08 | 0.94 | yes |
| 203 | 157.050 | C7H9O4+ | 4.89 | 7.00E-10 | 1.05 | yes |
| 204 | 157.086 | C8H13O3+ | | | | no |
| 205 | 157.122 | C9H17O2+ | | | | no |
| 206 | 157.159 | C10H21O+ | 5.09 | 5.77E-17 | 0.83 | yes |
| 207 | 158.154 | C9H20NO+ | 2.69 | 6.83E-15 | 0.89 | yes |
| 208 | 159.030 | C7H11S2+ | 3.02 | 2.68E-10 | | no |
| 209 | 159.065 | C7H11O4+ | | | | no |
| 210 | 159.102 | C8H15O3+ | | | | no |
| 211 | 159.138 | C9H19O2+ | | | | no |
| 212 | 161.045 | C7H13S2+ | 3.41 | 6.96E-11 | 1.01 | yes |
| 213 | 161.081 | C7H13O4+ | | | | no |
| 214 | 161.117 | C8H17O3+ | | | | no |
| 215 | 161.132 | C12H17+ | 2.36 | | 0.98 | yes |
| 216 | 161.154 | C9H21O2+ | 2.11 | | 0.91 | yes |
| 217 | 163.133 | C8H19O3+ | 4.68 | 3.20E-11 | 1.05 | yes |
| 218 | 169.195 | C12H25+ | 16.10 | 6.26E-22 | 1.03 | yes |
| 219 | 171.080 | C12H11O+ | 2.91 | 7.19E-12 | 0.95 | yes |
| 220 | 175.169 | C10H23O2+ | 4.30 | 7.32E-13 | 0.83 | yes |
| 221 | 177.164 | C13H21+ | 10.60 | 6.44E-19 | 0.76 | yes |
| 222 | 185.190 | C12H25O+ | 6.78 | 2.42E-20 | 0.91 | yes |
| 223 | 189.127 | C13H17O+ | 11.63 | 1.87E-14 | 1.05 | yes |
| 224 | 195.174 | C13H23O+ | 9.40 | 6.63E-19 | 0.73 | yes |
| 225 | 203.143 | C14H19O+ | 2.56 | 7.81E-06 | 1.02 | yes |
| 226 | 203.201 | C12H27O2+ | 5.42 | 2.36E-16 | 0.91 | yes |
| 227 | 205.195 | C15H25+ | 25.43 | 1.29E-23 | | no |
| 228 | 214.090 | C10H16NO2S+ | 3.94 | 2.44E-13 | 1.10 | yes |
| 229 | 223.133 | C13H19O3+ | 6.58 | 1.43E-09 | 1.07 | yes |
| 230 | 235.206 | C16H27O+ | 27.94 | 4.67E-13 | 0.89 | yes |
| 231 | 243.268 | C16H35O+ | 4.19 | 4.37E-07 | 1.05 | yes |
| 232 | 247.227 | C14H31O3+ | 4.53 | 1.71E-12 | 1.07 | yes |
| 233 | 259.206 | C18H27O+ | 3.75 | 2.75E-07 | 1.08 | yes |
| 234 | 277.216 | C18H29O2+ | 3.83 | 5.67E-07 | 1.09 | yes |

**Supplementary Table 2 | Compound identification (WA: water adduct ion, WL: water loss ion, AA: ammonia adduct ion)**

| IC map | Mass trace | Ion formula | Ion species | Identification | Ion name (GCxGC-QTOF isomer contributions) |
|---|---|---|---|---|---|
| 2 | 163.133 | C8H19O3+ | [M+H3O]+ | High corr. (C8H17O2+) | Caprylic acid WA |
| | 145.122 | C8H17O2+ | [M+H]+ | Library (GCxGC-QTOF) | Caprylic acid (with trace amounts of methyl heptanoate & 1,3-Dioxolane, 4-methyl-2-(2-methylpropyl)-) |
| | 129.127 | C8H17O+ | [M+H]+ | Library (GCxGC-QTOF), high corr. (C8H17O2+) | Octanal (with minor amounts of 2-octanone, ¼ of octanal) |
| | 147.138 | C8H19O2+ | [M+H3O]+ | High corr. (C8H17O+) | Octanal WA |
| | 89.060 | C4H9O2+ | [M+H]+ | Library (GCxGC-QTOF) | Butyric acid (with trace amounts of ethyl acetate, 1-4-dioxane and methyl propionate |
| | 83.086 | C6H11+ | [M+H]+ | Library (GCxGC-QTOF) | Non-specific fragment |
| | 111.117 | C8H15+ | [M+H]+ | Library (GcxGC-QTOF), high corr. (C8H17O+) | Non-specific fragment |
| | 113.132 | C8H17+ | [M+H]+ | Library (GCxGC-QTOF) | Non-specific fragment (with minor amounts of 1-octene) |
| 6 | 61.028 | C2H5O2+ | [M+H]+ | Library (GCxGC-QTOF) | Acetic acid |
| | 79.039 | C2H7O3+ | [M+H3O]+ | High corr. (C2H5O2+) | Acetic acid WA |
| | 43.018 | C2H3O+ | [M+H]+ | High corr. (C2H5O2+) | Acetic acid WL |
| | 163.133 | C8H19O3+ | [M+H3O]+ | High corr. (C8H17O2+) | Caprylic acid WA |
| | 127.112 | C8H15O+ | [M+H]+ | Library (GCxGC-QTOF) | Sulcatone |
| | 157.159 | C10H21O+ | [M+H]+ | Library (GCxGC-QTOF) | Decanal (with minor amounts of 2-decanone, 1/10 of decanal) |
| | 175.169 | C10H23O2+ | [M+H3O]+ | High corr. (C10H21O+) | Decanal WA |
| | 195.174 | C13H23O+ | [M+H]+ | Library (GCxGC-QTOF) | Geranylacetone |
| 11 | 135.102 | C6H15O3+ | [M+H3O]+ | High corr. (C6H13O2+) | Caproic acid WA |
| 13 | 113.023 | C5H5O3+ | [M+H]+ | Library (GCxGC-QTOF) | Citraconic anhydride |
| | 131.034 | C5H7O4+ | [M+H3O]+ | High corr. (C5H5O3+) | Citraconic anhydride WA |
| 14 | 59.049 | C3H7O+ | [M+H]+ | Library (GCxGC-QTOF) | Acetone |
| | 77.060 | C3H9O2+ | [M+H3O]+ | High corr. (C3H7O+) | Acetone WA |
| | 76.076 | C3H10NO+ | [M+NH4]+ | High corr. (C3H7O+) | Acetone AA |
| | 60.021 | Unknown | Unknown | High corr. (C3H7O+) | Acetone artefact peak |

**Supplementary Table 3 | Compound properties (structure, common name, RN: reference number (CAS), molecular formula, exact mass, volatility (bp: boiling point, °C, ACD/labs), polarity (logP, ACD/labs), and odour)**

| | | | |
|---|---|---|---|
| 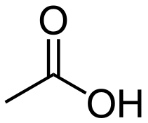 | Acetic acid, RN: 64-19-7<br>C2H4O2, 60.0211 u<br>bp: 117.1±3.0 °C, logP: -0.28<br>odour: vinegar | 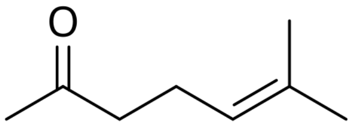 | Sulcatone, RN: 110-93-0<br>C8H14O, 126.1045 u<br>bp: 173.3±9.0 °C, logP: 2.09<br>odour: citrus-like |
| 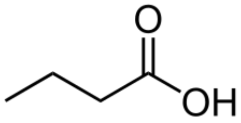 | Butyric acid, RN: 107-92-6<br>C4H8O2, 88.0524 u<br>bp: 164.3±3.0 °C, logP: 0.79<br>odour: vomit-like | 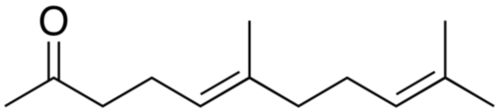 | Geranylacetone, RN: 3796-70-1<br>C13H22O, 194.1671 u<br>bp: 256 °C, logP: 4.13<br>odour: floral |
| 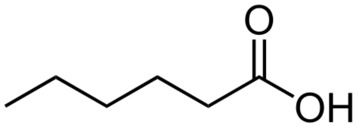 | Caproic acid, RN: 142-62-1<br>C6H12O2, 116.0837 u<br>bp: 204.6±3.0 °C, logP: 1.84<br>odour: goat-like | 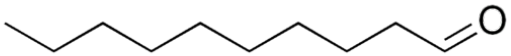 | Decanal, RN: 112-31-2<br>C10H20O, 156.1514 u<br>bp: 209.0±3.0 °C, logP: 4.09<br>odoru: fruity |
| 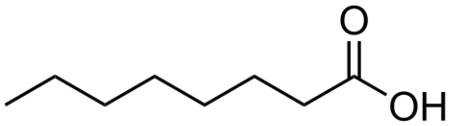 | Caprylic acid, RN: 124-07-2<br>C8H16O2, 144.1150 u<br>bp: 239.3±3.0 °C, logP: 2.90<br>odour: goat-like | 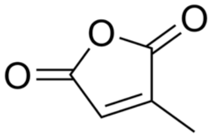 | Citraconic anhydride, RN: 616-02-4<br>C5H4O3, 112.0160 u<br>bp: 211.5±9.0 °C, logP: 0.23<br>odour: slightly sweet |
| 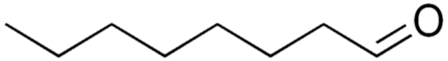 | Caprylic aldehyde, RN: 124-13-0<br>C8H16O, 128.1201 u<br>bp: 163.4 °C, logP: 3.03<br>odour: fruity | 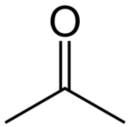 | Acetone, RN: 67-64-1<br>C3H6O, 58.0419 u<br>bp = 56.08±3.0 °C, log P: - 0.24<br>odor: pungent |

**Supplementary Table 4 | Subjective assessments of the VR scenario used for inducing fear, comparison between post neutral and post fear sessions**

State anxiety (STAI-Y1) and self-reported fear were assessed after the neutral and fear-induction scenarios using independent questionnaires (see Methods, "Experimental design and protocol", "Subjective assessments"). These measures were collected for all 45 participants enrolled in the experimental study. We compared post-neutral and post-fear scores using two-tailed paired Wilcoxon signed-rank tests, performed separately for STAI-Y1 and self-reported fear scores (Matlab signrank function). Both tests rejected the null hypothesis of a zero median paired difference, with higher STAI-Y1 and self-reported fear scores after the fear-induction scenario than after the neutral scenario. Full statistical results are reported below, with 95% confidence intervals computed around the Hodges–Lehmann estimate of the paired median difference. Boxplots of STAI-Y1 and self-reported fear scores are shown in Supplementary Fig. 1 and Supplementary Fig. 2, respectively.

| **Assessment** | **Z(45)** | **p-value** | **Median Post Neutral** | **Median Post Fear** | **Median averages of differences** | **95% CI** | **Effect size (r)** |
|---|---|---|---|---|---|---|---|
| **STAI-Y1** | -2.1643 | 3.04×10^(-2) | 37 | 42 | 4 | [0.5; 7.5] | 0.3226 |
| **Self-Reported Fear** | -4.5084 | 6.53×10^(-6) | 1 | 5 | 2.5 | [1.5; 3.0] | 0.6721 |

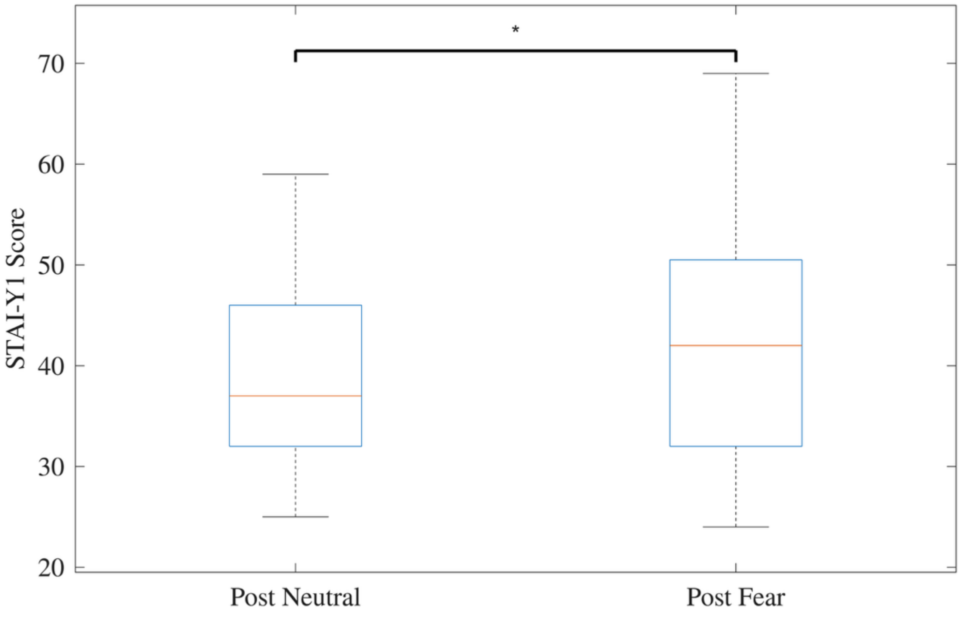


Supplementary Fig. 1: Boxplot of STAI-Y1 score measured after the Neutral and Fear sessions.

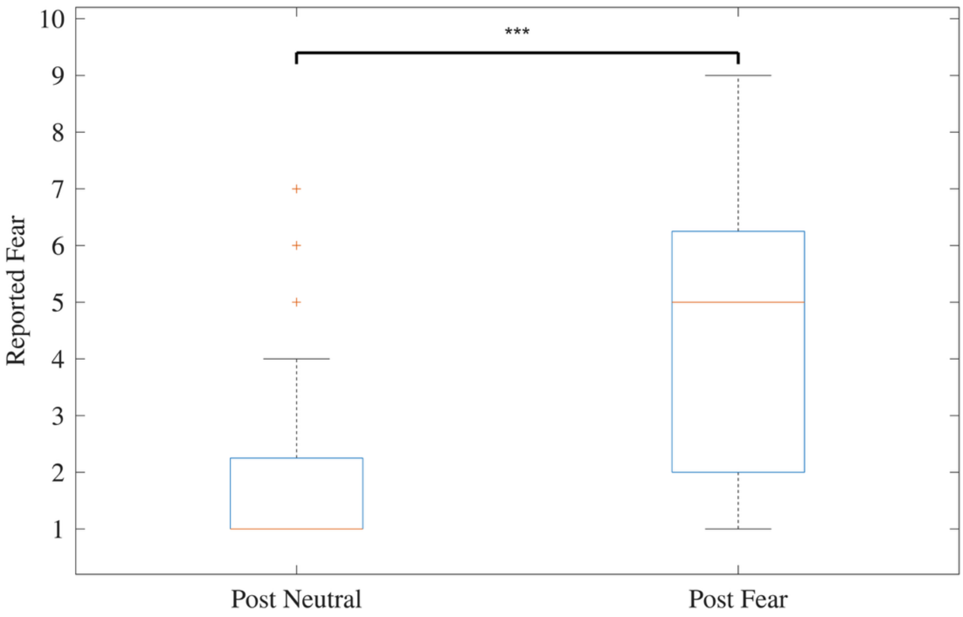


Supplementary Fig. 2: Boxplot of self reported fear score measured after the Neutral and Fear sessions.

**Supplementary Table 5 | Physiological assessment of the VR scenario used for inducing fear, comparison between neutral and fear sessions**

Electrodermal activity was recorded throughout the experimental session for the 45 participants enrolled in the study (see Methods, “Physiological data acquisition”). To assess whether the VR scenario elicited sympathetic activation, we computed EDASymp, a frequency-domain index of sympathetic activity defined as the power of the EDA signal in the 0.045–0.25 Hz frequency range [1]. One participant was excluded from this analysis because of EDA recording failure, leaving 44 participants with valid EDA data.

To compute EDASymp, EDA signals were first downsampled to 50 Hz and normalized by z-scoring. We then applied the cvxEDA decomposition algorithm [2] to estimate the skin conductance level (SCL) and skin conductance response (SCR) time courses (see Methods, “Data preprocessing and feature extraction”). The denoised EDA signal was obtained as the sum of SCL and SCR, separately for the neutral and fear-induction sessions. Power spectral density was estimated using Welch’s method with a 60-s Blackman window, 50% overlap and an FFT length equal to twice the window length. EDASymp was computed as the summed spectral power in the 0.045–0.25 Hz frequency band.

Neutral and fear-induction EDASymp values were compared using a two-tailed paired Wilcoxon signed-rank test (Matlab signrank function). The test rejected the null hypothesis of a zero median paired difference, with higher EDASymp during the fear-induction scenario than during the neutral scenario. Full statistical results are reported below, with 95% confidence intervals computed around the Hodges–Lehmann estimate of the paired median difference. EDASymp values across the two sessions are shown in Supplementary Fig. 3.

| Assessment | Z(44) | p-value | Median Neutral | Median Fear | Median averages of differences | 95% CI | Effect size (r) |
|---|---|---|---|---|---|---|---|
| **EDASymp** | -5.7651 | $8.16 \times 10^{-9}$ | -1.4651 | 2.8175 | 5.1609 | [3.6455; 6.6263] | 0.8691 |

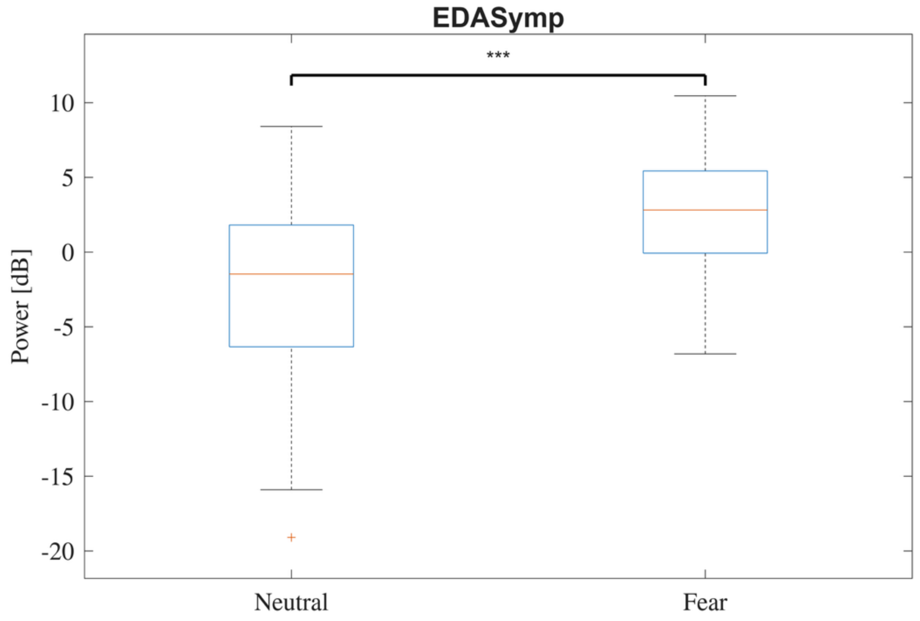


Supplementary Fig. 3: Boxplot of EDASymp measured during the Neutral and Fear sessions.
